# Mapping and identification of soft corona proteins at nanoparticles and their impact on cellular association

**DOI:** 10.1101/2020.02.05.924480

**Authors:** Hossein Mohammad-Beigi, Yuya Hayashi, Christina Moeslund Zeuthen, Hoda Eskandari, Carsten Scavenius, Kristian Juul-Madsen, Thomas Vorup-Jensen, Jan J. Enghild, Duncan S. Sutherland

**Author notes:** Corresponding author: Duncan S. Sutherland.

## Abstract

The current understanding of the biological identity that nanoparticles may acquire in a given biological milieu is mostly inferred from the hard component of the protein corona (HC). The composition of soft corona (SC) proteins and their biological relevance have remained elusive due to the lack of analytical separation methods. Here, we identified a set of specific corona proteins with weak interactions at silica and polystyrene nanoparticles by using an in situ click-chemistry reaction. We show that these SC proteins are present also in the HC, but are specifically enriched after the capture, suggesting that the main distinction between HC and SC is the differential binding strength of the same proteins. Interestingly, the weakly interacting proteins in the SC are revealed as modulators of nanoparticle-cell association, in spite of their short residence time. We therefore highlight that weak interactions of proteins at nanoparticles should be considered when evaluating nano-bio interfaces.

---

Nanoparticles (NPs) are promising agents for drug delivery and visualisation in vivo. Upon exposure to biofluids, the NPs acquire a ‘protein corona’ due to the adherence of host proteins on the NP surface. The composition of the corona is dependent on the types of nanoparticles and the biological sources ^1–3^ and considered to provide the NPs with a ‘biological’ identity ^4^ that affects stability, circulation time and cellular uptake/interactions and therefore has strong impact on the functional role for the NP ^5–10^.

Since the introduction of the biomolecular corona concept for NPs ^11,12^, identification of the proteins forming the corona has become an active research topic aiming to understand the particokinetics, cellular interactions, and mechanisms of toxicity^13^. In a complicated and dynamic process, proteins competitively adhere to the surface of nanoparticles and form a “Hard” (HC) and “Soft” corona (SC). HC proteins with a high binding affinity and low dissociation rate remain tightly bound to the surface, whereas SC proteins with a high dissociation rate are rapidly exchanged. At the surface of nanoparticles, proteins can undergo reorientation and conformational changes ^14,15^, presumably leading to at least a partially denatured state that has a reduced dissociation rate in a process referred to as “hardening”^16^. The evolution and dynamics of HC formation are relatively well studied ^11,17–19^; HC is established rapidly and the evolution of HC over time is only quantitative ^31^ with altered relative amounts, rather than the changes in protein composition expected from the Vroman effect ^20^. The current understanding is that the HC proteins - with their long residence time - give the nanoparticles a biological identity by presenting receptor-binding sites for cellular interactions with a biologically relevant time-scale ^21^. As SC proteins by definition have a shorter residence time on nanoparticles than HC proteins and making them difficult to isolate from free proteins of the mother liquid, their potential biological impacts have often been ignored. Recent work has developed approaches to quantify SC protein binding and address the potential of soft interactions to modulate toxicity by localized sulphidation at the surface of silver nanoparticles ^13^. However, so far no method has been reported for profiling SC proteins and several key open questions remain, including whether SC proteins are different from HC proteins and if there is a role for SC proteins in determining cellular interactions.

To address these critical questions, a comprehensive picture of corona composition and residence time for SC proteins is needed. Here, by developing an experimental approach based on click chemistry, we captured weakly interacting proteins along with HC proteins for mass spectrometry-based compositional profiling. We find that the majority of the SC proteins are not unique to SC but are also present in the HC representing different binding strength states of the same proteins. On the contrary, only a minor fraction of SC proteins were identified exclusively in the SC. Moreover, as our method forces SC proteins to stay in place by crosslinking, such that the SC proteins acquire time long enough for biological interactions, we were able to demonstrate a role for the SC proteins in cell association of nanoparticles, that was dependent both on the type of cells and nanoparticles.

## A novel click chemistry method captures SC proteins

Recently, the catalyst-free strain-promoted alkyne azide cycloaddition (SPAAC) “click” chemistry has gained interest in many biological and medical applications due to its high speed, efficiency, specificity, and bioorthogonality ^22–25^. Therefore, we have developed a SPAAC click reaction between azide-modified HC proteins on nanoparticles (HC-N_3_) and Dibenzocyclooctyne (DBCO)-activated SC proteins (FBS-D) (Fig.1a) in order to trap the transiently binding SC proteins on the NP surface (HC+SC sample). Silica nanoparticles (SNPs, 70 nm) and carboxylate-modified polystyrene nanoparticles (PsNPs, 100 nm) were used in this study as model nanoparticles ^26–28^. We used four control samples representing HC and FBS with and without chemical modifications (hard corona (HC), hard corona modified with azide (HC-N_3_), FBS-D added to HC (D Ctrl), FBS added to HC-N3 (N_3_ Ctrl)) and one HC+SC sample that encompasses proteins in both HC and captured SC states.

**Fig. 1.**
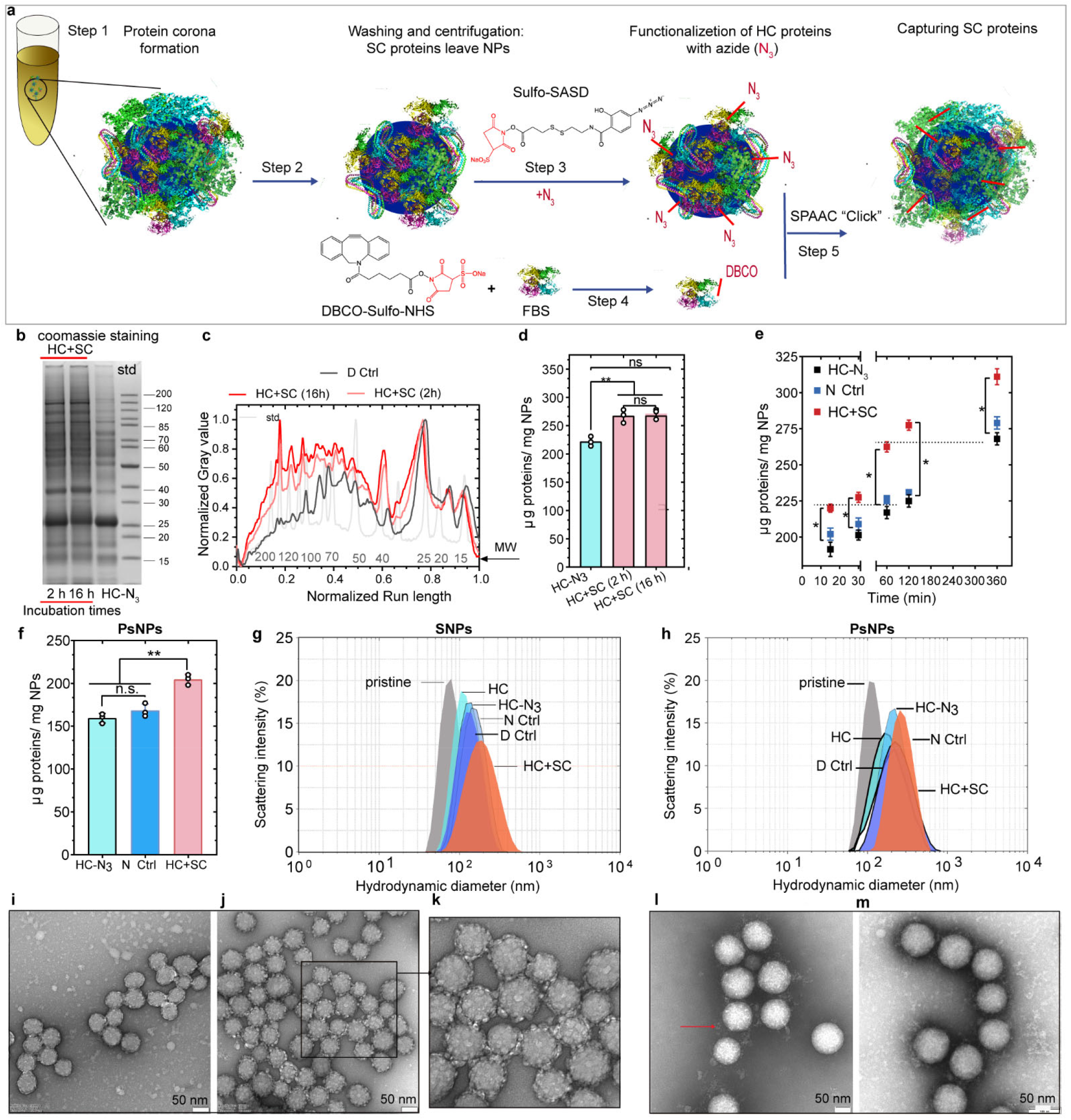
SPAAC click chemistry reaction and characterization of nanoparticle-corona complexes. **a,** Schematic representation of capturing SC proteins. After protein corona formation (steps 1 and 2), the HC proteins were modified with N_3_ by reacting with Sulfo-SASD (step 3) followed by a SPAAC “click” reaction (step 5) with FBS-D proteins (prepared in step 4). **b-d,** Effect of exposure time periods (2 and 16 h) in the click reaction evaluated by coomassie staining images (**b**) and densitometry analysis of SDS-PAGE gel (**c**), and quantification (**d**) of protein corona recovered from SNPs. SDS-PAGE analysis revealed a change in protein patterns after the click reaction, in particular, a characteristic increase in the 40 kDa bands and a higher intensity of bands between 50-200 kDa. Quantification data represented as the mean ± sd. of three independent experiments (n=3). **e,** Quantification of HC+SC proteins captured by click reaction on HC proteins formed on SNPs over different incubation times (15 min, 30 min, 1 h, 2h, 6 h). The SDS-PAGE image and densitometry analysis of the proteins are shown in Supplementary Fig. S4. **f,** Quantification of HC+SC proteins captured by click reaction on PsNPs. **g**,**h**, Hydrodynamic analysis of nanoparticle-corona complexes, SNPs (**g**) and PsNPs (**h**). **i-m**, Transmission electron microscopy (TEM) analysis of the SNPs@HC (**i**), SNPs@HC+SC (**j** and **k**), PsNPs@HC (**l**) and PsNPs@HC+SC (**m**). Scale bar, 50 nm. nomenclature: FBS-D: FBS proteins modified with DBCO, pristine silica nanoparticles (SNPs), pristine polystyrene nanoparticles (PsNPs), hard corona (HC), hard corona modified with azide (HC-N_3_), FBS-D added to HC (D Ctrl), FBS added to HC-N3 (N_3_ Ctrl), FBS-D added to HC-N_3_ (HC+SC).

We first determined the extent to which the click reaction could capture SC proteins (optimization of the click reaction is explained in the supplementary results). SDS-PAGE analysis revealed a change in protein patterns after the click reaction which was further validated by capturing fluorescently labelled proteins (Supplementary Fig. S3). Quantification of total protein per nanoparticle further confirmed the increase in the mass of the corona proteins (~50 μg/mg nanoparticles) after the click reaction. Extending the reaction time from 2 h to 16 h did not result in a further increase in mass (Fig.1d). The amount of SC proteins captured was positively correlated with the amount of HC proteins bound to the nanoparticles, which increased as a function of incubation time (Fig. 1e and Supplementary Fig. S4). Illustrating the applicability of this method to other types of nanoparticles, we observed comparable results for PsNPs (Fig. 1f and Supplementary Fig. S5).

Our click chemistry approach to capturing SC proteins maintained a colloidally stable population of nanoparticle-corona complexes with slightly increased hydrodynamic size and an increased particle heterogeneity (Fig. 1g-m and Supplementary Table 2). We further analyzed the formation of the protein corona by negative staining (TEM), which revealed a globular appearance of dehydrated proteins on SNPs (Fig.1 i-k and Supplementary Fig. S6), while a more diffuse appearance was observed for PsNPs (Fig.1l-m and Supplementary Fig. S7). Image analysis of the SNPs confirmed a broadened distribution of maximum Feret particle diameters with an increase in the mean size from 72 nm (HC) to 87 nm (HC+SC) (Supplementary Fig. S6d).

## SC is composed mainly of proteins already present in the HC, representing different binding states of the same proteins

Using the click reaction to fix the weakly interacting proteins in place, we were able to isolate SC proteins along with HC proteins by centrifugation and subject them to proteomic quantification by tandem mass spectrometry (LC-MS/MS).

A cluster of proteins specifically enriched in the HC+SC sample (based on the copy number of corona proteins per nanoparticle, Supplementary Information for details) was identified using bottom-up cluster analysis to construct two-way dendrograms along with a heatmap (Fig. 2a-f). For SNPs, 20 proteins were considered as SC proteins among the total of 80 proteins identified by LC-MS/MS, and only 4 out of the 20 SC proteins were uniquely captured after the click reaction (i.e. undetected in all HC controls), while the others were found in the HC controls to some extent (Fig. 2b). The total copy number of all proteins per NP increased 1.15-fold after the click reaction (cf. ~1.2-fold increase in the total protein mass in Fig. 1e), and the increase was mainly due to higher copy numbers of proteins belonging to the SC cluster (Fig. 2c) (e.g. 5.7-fold increase in the abundance of APO H; Table 1, “SNPs” and Fig. 2b). Interestingly, the absence of highly abundant serum proteins such as albumin in the SC cluster shows that our click chemistry method used in a competitive situation captures only proteins which are resident at the surface.

**Fig. 2.**
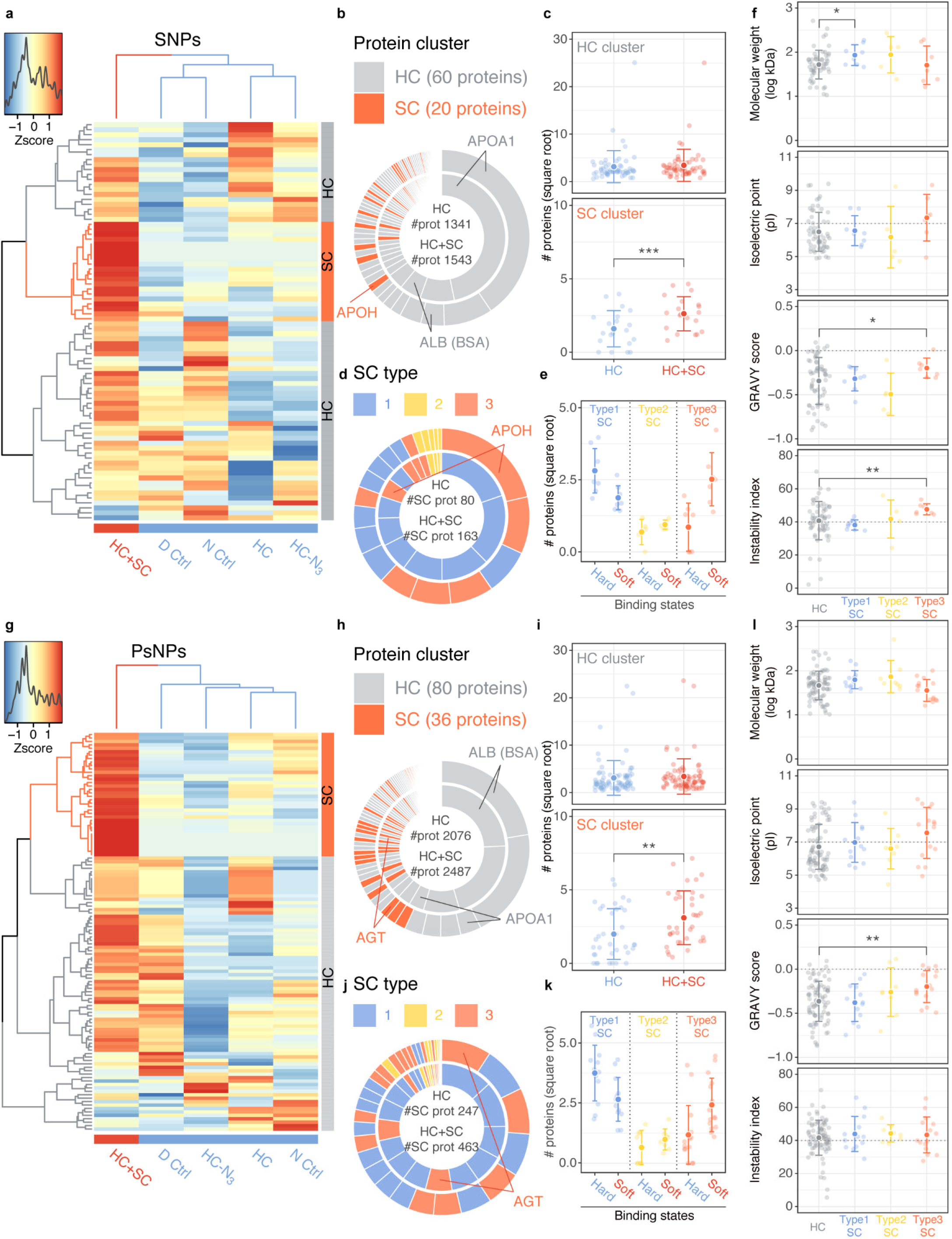
Identification of soft corona proteins and their classification according to the relative abundance in two binding states. **a,g,** A heatmap with two-way unsupervised hierarchical clustering analysis (UHCA) of the relative abundance of corona proteins recovered from SNPs (**a**) and PsNPs (**g**). Each row, a protein; each column, a protein corona sample. The number of proteins per nanoparticle is scaled to derive a z-score representing the relative abundance of each protein between the samples. A colour key along with the z-score distribution is depicted to the top left. Red and blue correspond to the number of proteins higher and lower than the average across all samples, respectively. The column dendrogram clearly separates the HC+SC sample from the rest. The row dendrogram reveals a putative SC cluster (coloured in orange) characterized by specific enrichment of the proteins in HC+SC. The complete heatmaps with the name of proteins are shown in Supplementary Fig. 8 and 9. **b,h,** The relative contribution of HC and SC cluster proteins to the total number of proteins (#prot) in HC (averaged from all four control samples) and HC+SC on SNPs (**b**) and PsNPs (**h**). The two doughnut charts represent the number percentages of each protein in HC (inner) and HC+SC (outer) where the SC proteins are coloured in orange. Proteins of particular interest are annotated. Both the number of different proteins and the total number of all proteins (#prot) per nanoparticle were larger in PsNPs than in SNPs. **c,i,** The square root number of proteins per nanoparticle for each protein from HC and SC clusters in SNPs (**c**) and PsNPs (**i**). While proteins from the HC cluster had the same quantity in HC and HC+SC samples, a significant increase was observed in the HC+SC sample for the proteins from the SC cluster, indicating the specific enrichment of these SC proteins. **d,j,** The relative contribution of three types of SC proteins to the total number of SC proteins (#SCprot) in HC (averaged from all four control samples) and HC+SC on SNPs (**d**) and PsNPs (**j**). The two doughnut charts represent the number percentages of each SC protein in HC (inner) and HC+SC (outer) where the three types of SC proteins are differentially coloured. Proteins of particular interest are annotated. In HC samples, the relative contribution of Type 1 SC proteins (blue) is larger than Type 3 SC proteins (orange), and vice versa in HC+SC samples. **e,k,** The square root number of proteins per nanoparticle for the two binding states (soft and hard) of SC proteins characterizing the three different SC types in SNPs (**e**) and PsNPs (**k**). The two binding states are assumed from the default presence of SC proteins in HC (hard binding) and upon “click” capturing of additional SC proteins in HC+SC (soft binding). Type 1 SC proteins are found more in the hard binding state and less in the soft binding state. Type 3 SC proteins have the opposite pattern. Type 2 SC proteins are intermediate. **f,l,** Four protein parameters (molecular weight, isoelectric point, GRAVY score, and instability index) are compared between the HC proteins and each type of SC proteins. Type 3 SC proteins show a significantly higher GRAVY score and instability index on SNPs (**f**) and higher GRAVY score on PsNPs (**l**). The comparison of the number-weighted average of the four parameters characterizing the overall protein property of the corona is shown in Supplementary Fig.11 **c,e,f,i,k,l,** Values are shown for individual proteins (soft coloured) and the mean ± sd. (solid coloured). Where relevant, Student’s *t*-test (**c,i**) or Welch’s unequal variance *t*-test (**f,l**) were performed. * *p*<0.05; ** *p*< 0.01; *** *p*<0.001.

**Table 1.** 20 most abundant proteins in HC and HC+SC samples and 20 most abundant SC proteins on SNPs and PsNP.

| SNPs |  |  |  |  |
| --- | --- | --- | --- | --- |
| No | 20 most abundant proteins in HC samples | 20 most abundant proteins in HC+SC samples | 20 most abundant proteins in the soft binding state | fold increase |
| 1 | APO-AI/46 % | APO-AI/40.5 % | APO H | 5.71 |
| 2 | Hemoglobin fetal subunit beta/8.7% | Hemoglobin fetal subunit beta/8.9 % | Complement C4 | 6.22 |
| 3 | Serum albumin/4.3 % | Serum albumin/4.3 % | UP2 | new |
| 4 | Hemoglobin subunit alpha/4 % | Hemoglobin subunit alpha/ 4 % | UP3 | new |
| 5 | Alpha-1-antiproteinase | Alpha-1-antiproteinase | UP1 | 3.31 |
| 6 | Kininogen-1 | Tetranectin | Cystatin-C | new |
| 7 | Plasminogen | Alpha-2-HS-glycoprotein | Fibulin-1 | 2.80 |
| 8 | Histidine-rich glycoprotein | Plasminogen | Collagen alpha-2(I) chain | 4.09 |
| 9 | Alpha-2-HS-glycoprotein | APO H | SERPINA11 protein | new |
| 10 | Apolipoprotein E | Serotransferrin | Fibronectin | 2.36 |
| 11 | Antithrombin-III | Histidine-rich glycoprotein | Alpha-S1-casein | 1.96 |
| 12 | Alpha-fetoprotein | Kininogen-1 | Thrombospondin-1 | 1.81 |
| 13 | Serotransferrin | Antithrombin-III | Complement C3 | 1.87 |
| 14 | Alpha-2-macroglobulin | Apolipoprotein E | Inter-alpha-trypsin inhibitor H3 | 1.60 |
| 15 | Protein S100-A12 | Alpha-2-macroglobulin | Inter-alpha-trypsin inhibitor H2 | 1.71 |
| 16 | Kininogen-2 | Alpha-fetoprotein | Hemopexin | 1.61 |
| 17 | Hemoglobin subunit beta | Insulin-like growth factor-binding 2 | Pigment epithelium-derived factor | 1.48 |
| 18 | Actin, cytoplasmic 1 | Apolipoprotein M | Prothrombin | 1.22 |
| 19 | Prothrombin | Complement C3 | Antithrombin-III | 1.32 |
| 20 | Apolipoprotein M | Hemoglobin subunit beta | Alpha-2-macroglobulin | 1.25 |
| PsNPs |  |  |  |  |
| No | 20 most abundant proteins in HC samples | 20 most abundant proteins in HC+SC samples | 20 most abundant proteins in the soft binding state | fold increase |
| 1 | Serum albumin/ 24.2 % | Serum albumin/22.3 % | Angiotensinogen | 2.19 |
| 2 | Hemoglobin fetal subunit beta/ 20.3% | Hemoglobin fetal subunit beta/20.3 % | Alpha-1-antiproteinase | 1.57 |
| 3 | Hemoglobin subunit alpha/3.8 % | APO-AI/3.8 % | Serpin family G member 1 | 2.35 |
| 4 | APO-AI/9.2 % | Hemoglobin subunit alpha/3.7 % | Serotransferrin | 1.59 |
| 5 | Plasminogen | Plasminogen | Tetranectin | 9.47 |
| 6 | Alpha-fetoprotein | Alpha-fetoprotein | Plasma serine protease inhibitor | 1.61 |
| 7 | Alpha-2-HS-glycoprotein | Alpha-1-antiproteinase | Cystatin-C | 22.22 |
| 8 | Alpha-1-antiproteinase | Serotransferrin | Complement factor I | 23.39 |
| 9 | Kininogen-1 | Angiotensinogen | Alpha-2-macroglobulin | 1.59 |
| 10 | Serotransferrin | Kininogen-1 | Complement factor H | 1.67 |
| 11 | Apolipoprotein E | Antithrombin-III | Insulin-like growth factor-binding 2 | 1.55 |
| 12 | Beta-2-glycoprotein 1 | Plasma serine protease inhibitor | Plasma kallikrein | 1.51 |
| 13 | Alpha-S1-casein | APO H | Secreted phosphoprotein 24 | 2.66 |
| 14 | Plasma serine protease inhibitor | Serpin family G member 1 | Fetuin-B | 1.33 |
| 15 | Kininogen-2 | Alpha-2-HS-glycoprotein | Hemopexin | 4.93 |
| 16 | Antithrombin-III | Sulfhydryl oxidase | Coagulation factor XII | 1.81 |
| 17 | Fetuin-B | Apolipoprotein E | Kininogen-2 | 1.25 |
| 18 | Actin, cytoplasmic 1 | Fetuin-B | Protein AMBP | 1.28 |
| 19 | Angiotensinogen | Kininogen-2 | Ig-like domain-containing protein | 11.98 |
| 20 | Beta-lactoglobulin | Alpha-2-macroglobulin | Fibulin-1 | 5.34 |

Proteins highlighted in the green are the proteins among the 20 most abundant proteins that appear to be in both HC and SC binding states. \* defines the SC proteins that are present on both SNPs and PsNPs.

Most of the SC proteins were also found in the HC, leading us to rethink our initial hypothesis that the SC is formed from different proteins from those in the HC. Our results rather indicate that the same proteins could have different binding constants and that SC proteins are those capable of both stable and transient interactions. For simplicity, we describe the SC proteins as generally having two binding states: “hard” and “soft”. Accordingly, we classified SC proteins into three types based on the relative copy numbers in the hard versus soft binding state: Type-1 SC proteins have more copies undergoing hard interactions, Type-2 SC proteins have similar copy numbers in the HC and SC, and Type-3 SC proteins have more copies undergoing soft interactions (2d-e). Of particular note is that there was a tendency for Type-3 SC proteins to have a higher GRAVY score and instability index, suggesting that these proteins are inherently less hydrophilic and less stable in serum(Fig. 2f). Neither the isoelectric point (Fig. 2f) nor multiparametric combinations of the four protein parameters revealed particular trends for the Type-3 SC proteins (Supplementary Fig. S10). The number-weighted averages of the overall protein characteristics of the HC controls and the HC+SC sample did not show a particular propensity indicating that the click reaction itself does not preferentially capture proteins with a specific parameter (Supplementary Fig. S11). For SNPs, the same set of experiments was performed using another type of stripping buffer described previously for LC-MS/MS ^29^ and we observed a similar pattern of SC proteins captured via the click chemistry approach (data not shown).

To look for a possible particle type-dependency, we applied the same method to analyze SC proteins on PsNPs. This revealed a slightly higher SC protein mass (1.2-fold increase after the click reaction) and more individual SC cluster proteins (36) than for SNPs (Fig. 2g-i). Notably, while there was a 30% overlap of HC cluster proteins between SNPs and PsNPs, only 7% of SC cluster proteins were common to both nanoparticle types (Supplementary Fig. S12). Type 3 SC proteins from PsNPS did not share properties except for the GRAVY score that tended to have a higher value (less hydrophilic) than HC cluster proteins, as also observed for the SNPs (Fig. 2l, Supplementary Figs. S10 and S11).

After the click reaction, the theoretical coverage ratio of NPs by corona proteins increased from 0.80-1.01 to 0.97-1.25 (for SNPs) and from 0.70-0.99 to 0.93-1.20 (for PsNPs), depending on the orientation of proteins on the NPs. It should be noted, however, that our click chemistry approach relies on the presence of an azide group on an HC protein within the reach (~1.8 nm) of the DBCO moiety of SC proteins, thus likely underestimating the amount of SC proteins, particularly for low HC coverages. Nevertheless, the derived coverage ratio is consistent with previous studies claiming that the corona formed from serum consists essentially of a monolayer ^2,30^. On this basis, we hypothesize that both HC and SC proteins co-exist within a loosely defined monolayer covering the nanoparticle surface, rather than in separate layers, and that SC proteins may dissociate from the surface during centrifugation. This fits well with our two binding states model introduced above and suggests that the monolayer of proteins becomes less dense as SC proteins dissociate, partially exposing the bare surface of the nanoparticle.

## APO H shows multiple binding strengths and outcompetes BSA for binding at HC coated SNPs

Apolipoprotein H (APO H) is among the most enriched proteins in Type 3, representing the major SC proteins on SNPs but not on PsNPs (Fig. 2 and Table 1). To test the ability of APO H to bind SNPs in a ‘hard’ and ‘soft’ state, a series of competition studies were performed using BSA and FBS. BSA was selected as the direct competitor since it is an HC cluster protein and has a higher affinity for the surface of SNPs than APO H. We started with uncoated SNPs and confirmed that APO H shows weaker binding to SNPs in the presence of the competitor BSA (Fig.3 a,b). We then let the HC pre-assemble on SNPs and captured APO H’s soft interactions through the click chemistry approach. Unlike the simple race for the bare surface, APO H was effectively enriched by the click reaction, outcompeting BSA with negligible influence of the concentration ratio between the two (Fig. 3 c,d). The low competitiveness of BSA for soft interactions with SNPs was also supported by the reduced capture of BSA via the click chemistry reaction, despite its high abundance, in the presence of FBS (Fig.3 e,f). Interestingly, in a parallel competition study the disease-related α-Synuclein (α-Syn) showed weak but specific binding to the HC proteins on SNPs unaffected by the presence of FBS proteins (Supplementary Fig. S14).

**Fig. 3.**
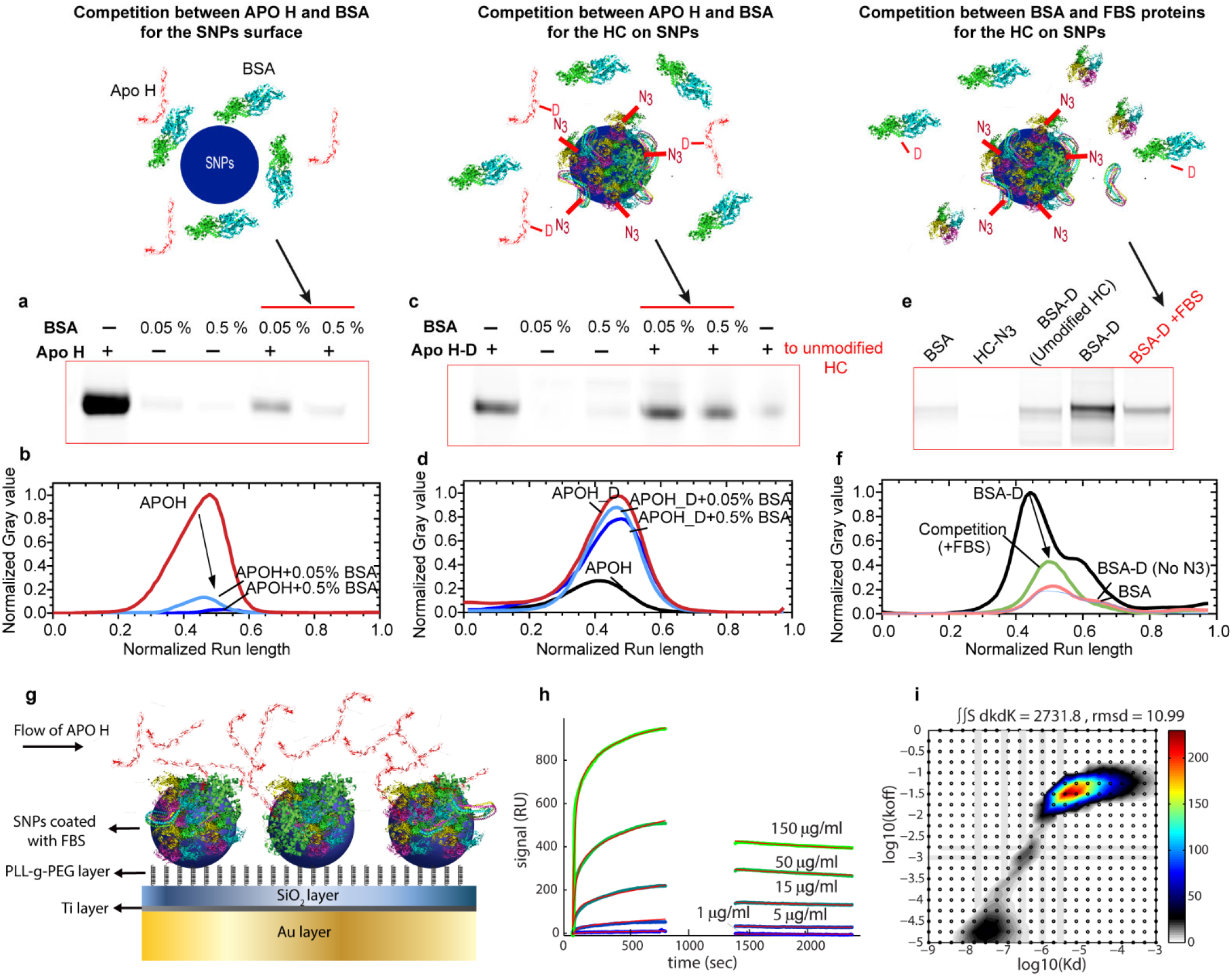
CY5 labeled APO H and BSA competition for the surface of SNPs and HC. **a-d** fluorescence images and densitometry analysis of cut-out SDS-PAGE gels of the addition of 30 μg/ml of APO H-CY5 protein corona on SNPs (**a,b**) and 30 μg/ml of APOH-CY5-DBCO on HC_N3 on SNPs (**c,d**) formed in the presence and absence of either 0.05 % or 0.5 % BSA. APO H-CY5-DBSO was added to SNPs@HC without N3 as a control sample in (**b**). **e,f,** The competition study between BSA and other FBS proteins for SNPs@HC-N3. fluorescence image of a cut-out SDS-PAGE gel of proteins recovered from the corona formed on SNPs (**e**) and its densitometry analysis (**f**). In this experiment, BSA-CY5-DBCO alone or spiked with FBS proteins was added to SNPs@HC-N3. The addition of BSA-CY5 to HC-N3 and BSA-CY5-DBCO to HC without N_3_ were done as controls. The complete SDS-PAGE gels are shown in the supplementary Fig. 13. **g-i** Surface Plasmon Resonance (SPR) characterization of APO H interaction with SNPS@HC. The schematic representation of the SPR characterization is shown in (**g**). First, SNPs were immobilized on a protein resistant polymer, PLL-g-PEG, creating an array of SNPs on a protein resistant background. Then, protein corona was formed by injecting 1 % FBS onto the immobilized NPs (shown in supplementary Fig.15). Subsequently, APO H was injected in different concentrations (1, 5, 15, 50, and 150 μg/ml) (**h**). Two-dimensional fits were applied to the data to calculate K_d_ and K_off_ values for different populations of APO H binding to the SNPs@HC (**i**). For the fitting, data from around the rinsing was omitted, due to too few data points. The complete data of two technical repeats are shown in supplementary Fig.15.

To study the soft binding property of APO H without click reactions, we utilised an immobilised-nanoparticle SPR setup to quantify the interaction of APO H with FBS proteins deposited onto SNPs ^31^ (Fig. 3g). When APO H was added to SNPs pre-coated with 1% FBS, a fraction of APO H remained associated with SNPs in a concentration-dependent manner (Fig. 3h). K_d_ and k_off_ values (supplementary Table S3) were obtained by fitting a two dimensional equation to the APO H data (Fig.3i), which shows a major population (88 ± 4.4 % of the signal) with high K_d_ values and fast off-rate and a small smeared population (9.5 ± 3.5 % of the signal) with low K_d_ values and slow off-rate. The fraction of APO H being removed at rinsing is also considered as another population with even lower binding strength. This study confirms the differential binding strength of APO H, where the major and minor fractions have high and low dissociation rates, representing soft and hard states of interactions on SNPs, respectively.

## SC proteins modulate cell association of nanoparticle-protein complexes in a cell- and particle type-dependent manner

Conceptually, pristine nanoparticles spontaneously bind to cell membranes in a non-specific manner in serum-free conditions, while protein coronas, in general, reduce this non-specific interaction by blocking the surface ^21^ and potentially increase specific interactions. For cell association (CA) via specific interactions, significant focus has been given to the HC proteins that have a residence time sufficient to support biological interactions (e.g. receptor ligation) ^19,32,33^. To elucidate the contribution of SC on CA, seen as the net effect of non-specific interactions of cells with the particle surface accessible by the dynamic SC proteins and specific interactions with the HC and SC (Supplementary Fig.S18), the click chemistry method was employed, which harden and transform the SC proteins to HC providing them a biologically relevant residence time. Phagocytic differentiated macrophage-like THP-1 cells (THP-1 macrophages), which express a family of cell surface recognition receptors ^34^, and human brain cerebral microvascular endothelial cells (hCMEC/D3), as a non-phagocytic counterpart were used in this study.

A preliminary study showed that cells readily interact non-specifically with bare particle surfaces of pristine nanoparticles which in general decreases from serum-free (SF) conditions to BSA media and then decreases further in FBS media, more significant for SNPs (Supplementary Fig. S19 and Fig. 4a). The effect of FBS on CA for the two particle types (over time of HC formation) was markedly different (Fig 4a-d) suggesting some involvement of specific interactions for SNPs. PsNPs, where BSA was the dominating protein in the corona, showed a systematic reduction of CA as the HC formed for both cell types, more significant for late corona (2 and 6 h). In contrast, SNPs, where APO-A1 was the major corona protein, showed an increased (for THP-1 cells) or unchanged (with hCMEC/D3 cells) CA as the HC formed, suggesting a different role for the dominant HC proteins in driving cell adhesion. Interestingly for SNPs, an opposite trend is seen when FBS-formed HC is studied in BSA media (with a pattern more similar to PsNPs).

**Fig. 4.**
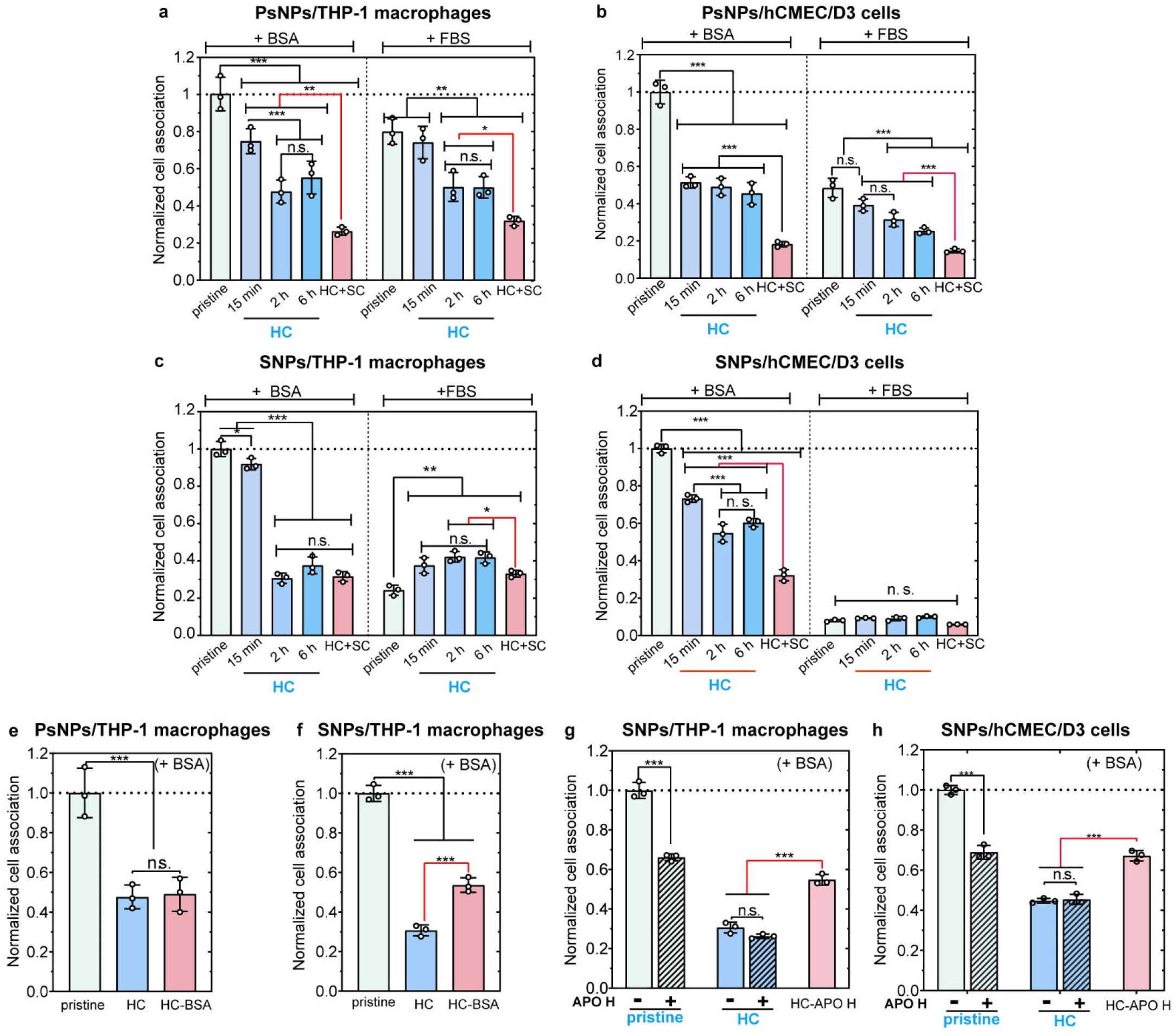
Cell association of nanoparticle-corona complexes in phagocytic and non-phagocytic cells. **a-d**, Flow cytometry was used to quantify the cell association of 50 μg/ml of nanoparticle-corona complexes in THP-1 macrophages (PsNPs,**a** and SNPs, **c**) and hCMEC/D3 cells (PsNPs,**b** and SNPs, **d**). The cells were exposed to the pristine NPs, coated with HC formed over different FBS exposure times (15 min, 2 h, and 6 h), and with HC+SC, for four hours in RPMI containing BSA or FBS.**e,f,** THP-1 macrophages were exposed to PsNPs@HC-APO H (**e**) and SNPs@HC-APO H (**f**) for 4 h in RPMI (BSA is crosslinked on HC by using a click reaction). **g,h,** THP-1 macrophage cells (**g**) and hCMEC/D3 cells (**h**) were exposed to SNPs or SNPs@HC for 4 h in RPMI containing BSA with or without 30 μg/ml APO H. The cells were also exposed to SNPs@HC_APO H (APO H is crosslinked on HC by using a click reaction). The flow cytometry data were normalized to the pristine nanoparticles values in the RPMI supplemented with BSA. Bars show mean ± sd. of three independent experiments. * p<0.05; ** p< 0.01; *** p<0.001; n.s., not significant. The cell gating data which was used to identify single cells are shown in Supplementary Fig. 16.

The artificial hardening of the SC proteins (HC+SC) generally decreased the CA of PsNPs in both cell types and media conditions (FBS or BSA) compared to particles with HC and short-lived dynamic SC, while SNPs showed decreased CA in THP-1 macrophages in FBS medium and hCMEC/D3 cells in the BSA medium (Fig. 4a-d). No difference in the localization of nanoparticles in the cells was observed by confocal microscopy (Supplementary Fig. S17). The hardening was also confirmed by a reduced ability of proteins to leave NPs in a serum-free medium (Supplementary Fig. S20 a,b) or for proteins to be exchanged with medium proteins such as albumin (Supplementary Fig. S20 c), which may explain why nanoparticles with hardened HC were less affected by changing media than pristine nanoparticles. The artificial cross-linking gives a more significant effect on CA than hardening made through evolution from a loosely attached toward a largely irreversibly attached protein during incubation time ^35^, as even long formation times for HC do not reduce the amount of SC proteins detected (Fig. 1e)

To confirm if specific proteins in HC and SC can enhance cell adhesion, BSA was crosslinked onto HC on both SNPs and PsNPs and APO H onto HC on SNPs. Surface-bound BSA has been reported to bind scavenger receptors on macrophages ^19^ and BSA specifically binds to FcRn receptors in endothelial cells ^36,37^. While BSA-crosslinked SNPs increased the CA in THP-1 cells, there was no effect on BSA-linked PsNPs (Fig. 4e,f). The increase in the CA of SNPs with crosslinked BSA implies that surface-bound BSA is recognized by cell receptors, and since BSA is the main protein forming the protein corona of PsNPs, crosslinking BSA did not significantly change its already relatively high contribution. APO H as a major but weakly interacting protein in the SNP corona is described as an opsonin for the mononuclear phagocyte systems, which also binds phospholipids in membranes ^38,39^. Crosslinked APO H could increase the CA of SNPs in both phagocytic and non-phagocytic cell lines (Fig. 4g, h); however, the addition of APO H to BSA medium decreased CA of pristine nanoparticles and had no effect on CA of nanoparticles with HC.

We interpret the reduced CA by crosslinking of SC from FBS as indicating that the dynamic SC proteins keep regions of the particle surface free from HC and available for interaction with cells despite the formed protein corona, allowing non-specific binding of the nanoparticle surfaces to cell membranes, dependent on their surface chemistry. Crosslinking would then decrease both the possibility for corona proteins to be exchanged with medium proteins and to reveal empty patches on the nanoparticle surface and enable non-specific cellular association. Here we show that SC proteins can contribute non-specifically to CA by revealing bare NP surfaces while hardening of individual SC proteins can contribute to CA through specific interactions.

## Conclusion

We describe here a novel, general and simple capture process based on click chemistry, which enabled the identification of weakly interacting proteins along with the long-lived protein corona forming around nanoparticles in complex media. For two different particle types, we find that the majority of the captured proteins are not unique to SC but also present in the HC indicating that the same proteins can have both strong and weak interactions with nanoparticles or pre-adsorbed neighbouring proteins.

SC proteins can thus be classified into 3 types based on the relative distribution between the hard and soft binding states (Fig. 5a). We propose that neighbouring proteins can restrict the hardening of SC proteins sterically limiting the surface area available for protein unfolding. By artificially hardening the SC proteins, we were able to “turn off” their dynamic nature of dissociation revealing a role in cellular interactions as caretaker proteins that can both prevent irreversible blocking of the surface and also allow higher affinity interactions between cell membranes and transiently-bare nanoparticle surfaces (Fig. 5b). The potential for cell-NP surface interactions suggests that the properties of the bare particle surface can still directly influence CA even after a fully developed protein corona is formed.

**Fig. 5.**
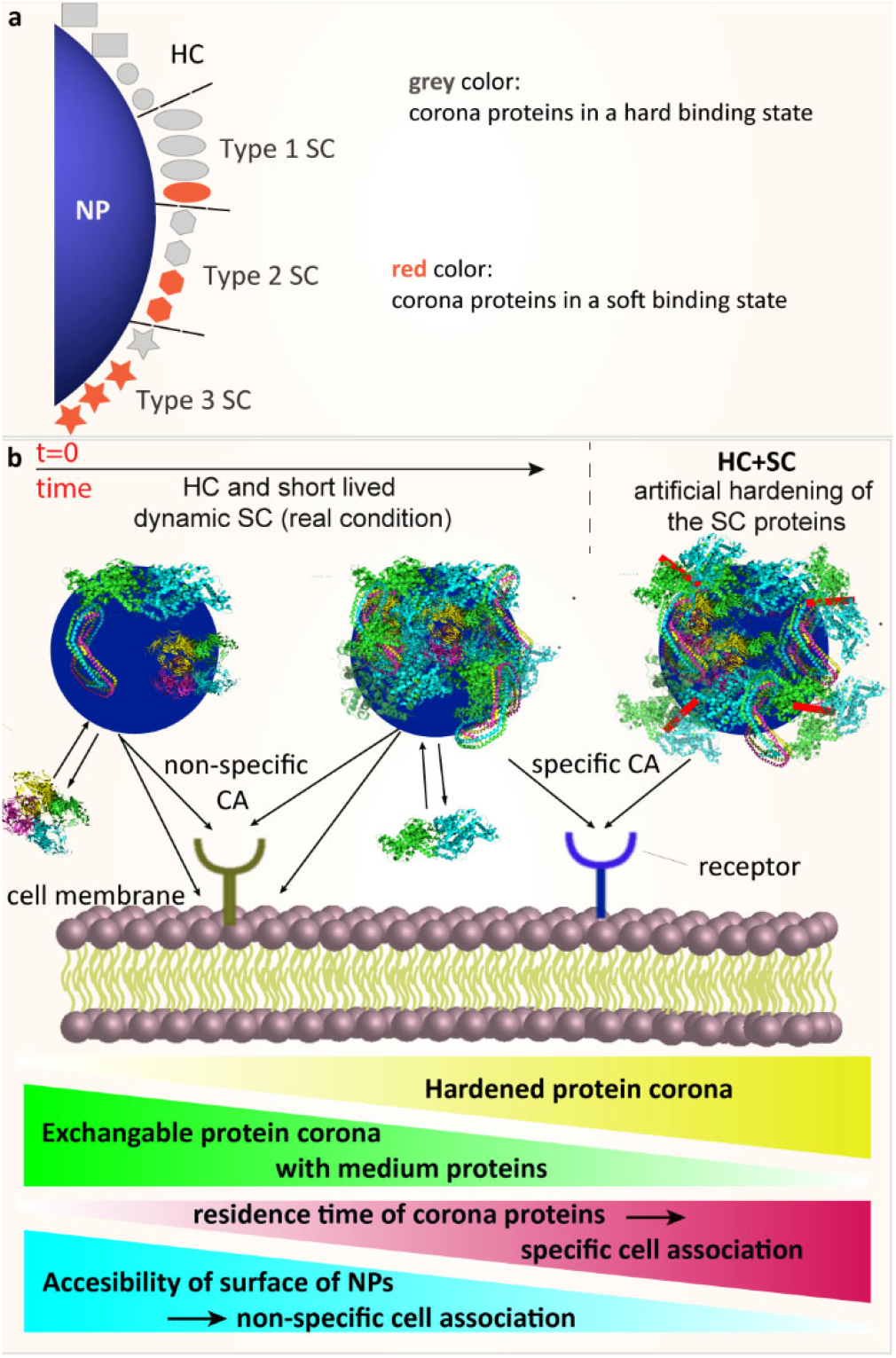
Schematic representation of the proposed model for the formation of hard and soft coronae and their contribution to the cell association of nanoparticle-corona complexes. **a,** Three types of SC proteins were identified by LC-MS/MS analysis after the click reaction. The SC proteins are mainly the same as HC proteins while the process of hardening does not define the protein corona composition but rather changes the binding strength of the already formed soft protein corona. **b,** Hardening of HC through either more incubation time or crosslinking SC by using a click reaction prepares a biological relevant residence time for corona proteins and reduces the exchange of corona proteins with medium proteins and accessibility of surface of NPs. The net effect of specific and non-specific associations will determine the fate of nanoparticles in contact with cells.nomenclature: CA: cell association

## Materials and Methods

All chemicals were of analytical grade and used as received.

### Nanoparticles

The 70 nm fluorescent plain silica nanoparticles (sicastar-greenF, fluorescein isothiocyanate-labeled; ex/em = 485/510 nm) was purchased from micromod Partikeltechnologie GmbH (Germany). The fluorescent carboxylate-modified polystyrene nanoparticles (FluoSpheres™ Carboxylate-Modified Microspheres, 0.1 μm, yellow-green fluorescent (505/515) was purchased from Thermo Fischer.

### Click-Chemistry reagents

Sulfo SASD (Sulphosuccinimidyl-2-(p-azidosalicylamido) ethyl-1,3-dithiopropionate) was purchased from G-biosciences. Dibenzocyclooctyne (DBCO)-Sulpho-NHS and DBCO-Sulpho-Cyanine5 were purchased form click chemistry tools and Genabioscience companies, respectively. Poly(L-lysine)-graft[3.5]-Poly(ethylene glycol)(2) (PLL-g-PEG) was purchased from SuSoS Ag.

### Fluorescent reagents

Sulpho-NHS-Cyanine5 was purchased form Lumiprobe. Hoechst 33342 Solution (20 mM) was purchased from Thermo Fischer. Phalloidin–Tetramethylrhodamine B isothiocyanate was purchased from Sigma-Aldrich.

### Cell media

Penicillin-Streptomycin, Gibco heat-inactivated Fetal Bovine Serum (FBS), Dulbecco’s Modified Eagle Medium (DMEM), and RPMI 1640 Media were purchased from Thermo Fischer. PMA Phorbol 12-myristate 13-acetate was purchased from Sigma-Aldrich.

### Nanoparticle incubation with FBS

FBS was first centrifuged at 16000 g for 3 min to remove any insoluble aggregates. Then, the protein supernatant and NPs solutions were pre-incubated at 37 °C for 10 min before mixing. The NPs were then exposed to FBS for different time points (15 min, 30 min, 1h, 2 h, and 6h) in darkness at 37 °C in Protein Lobind tubes.The ratio of total particle-surface area to FBS volume was kept constant for all nanoparticles. 0.4 mg SNPs (70 nm) and 0.3 mg PsNPs (100 nm) were used. The nanoparticle-corona complexes were isolated from unbound and loosely bound FBS proteins by centrifugation at 18000 g, 20 min. Pellets were washed three times with PBS by centrifugation at 20000 g, 20 min and resuspended in PBS for further analysis.

### Azide modification of hard corona (HC) proteins

To generate azide functionalized HC proteins, Sulpho-SASD at different concentrations (0, 0.018, 0.036, 0.09, 0.18, 0.35, 0.55, 1.8 mM) was added to the nanoparticles@HC at final concentration of 0.4 mg/ml. Sulpho-SASD contains a dithiol in the structure that can be cleaved by a reducing agent for electroporation analysis. The complex was incubated at 37 °C for 1 h to allow the reaction between the Sulpho-NHS and amine groups on HC proteins. The azide modified nanoparticle-corona complexes were separated from unreacted Sulpho-SASD by centrifugation at 20000 g, 20 min. The pellet was washed three times with PBS and then resuspended in PBS for further steps. To characterize the azide groups on HC proteins, a click reaction between the azide groups and DBCO-Sulpho-Cy5 was employed. By measuring the excited fluorescence (at 646 nm) of proteins at 664 nm, the minimum number of azide groups was calculated on HC proteins, assuming that each DBCO-Sulpho-CY5 reacts with one Sulpho-SASD. The number of HC corona proteins was measured by two methods: 1) by calculating the average molecular weight of proteins on SDS-PAGE and quantification of proteins by BCA assay, 2) LC-MS/MS analysis. In all steps, the number of nanoparticles was measured by reading the fluorescence of NPs.

### Modification of proteins with DBCO-Sulpho-NHS and DBCO-Sulpho-CY5

The reactive DBCO Sulpho-NHS at various final concentrations (0, 0.2, 0.4, 0.8 mM) was added to a tube containing 10 % FBS solution (4.2 mg/ml) at PBS (pH 7.4). The molar ratio of the crosslinkers to proteins (if we assume all proteins are BSA) at the selected concentrations is equal to 0, 2.5, 5 and 10. The system was then incubated at 37 °C for 1 h to allow the reaction between the Sulpho-NHS and amine groups on proteins. The reaction was quenched by Tris at a final concentration of 50 mM. The unreacted DBCO molecules were eliminated by a Sephadex G-25 in PD-10 Desalting Columns (GE Healthcare Life Sciences). To make fluorescently labeled DBCO modified FBS proteins or individual proteins (BSA or APO H), proteins were incubated with both DBCO-Sulpho-NHS and Sulpho-NHS-Cy5 (at the molar ratio of crosslinkers to proteins of 5). The degree of labelling (DOL) of proteins was calculated by measuring the UV spectrum of conjugates using the following equation 1:

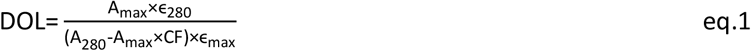

where A_max_ and A_280_ are absorbances of the conjugate solution measured at 280 nm and at λ_max_ of the crosslinker or dye, respectively. λ_max_ values for DBCO Sulpho-NHS and NHS-Sulpho-CY5 are 309 and 646 nm, respectively. ε_280_ and ε_max_ are the extinction coefficient of proteins (for FBS proteins, we used ε_280_ value of albumin, 66433 M^−1^cm^−1^) and DBCO Sulpho-NHS or NHS-Sulpho-CY5, whose values were taken as 12000 and 271000 M^−1^cm^−1^, respectively. CF is the correction factor of each crosslinker or dye which is required to eliminate the contribution of the dye at 280 nm. A_max_ and A_280_ are absorbances of the conjugate solution measured at 280 nm at λ_max_ of the crosslinker or dye.

### Crosslinking SC proteins on HC proteins

In order to capture weakly interacting proteins on nanoparticle-corona complexes, the azide modified particles were incubated with DBCO modified FBS (FBS-D) for 2 h at 37 °C. The azide modified nanoparticle-corona complexes were separated from free proteins by centrifugation at 20000 g, 20 min. The pellet was washed three times with PBS and then resuspended in PBS for further steps.

### Characterization of nanoparticle–corona complexes

Nanoparticles were characterized by transmission electron microscopy (TEM), dynamic light scattering (DLS); and zeta potential measurements. Both zeta potential and hydrodynamic diameter of nanoparticles were measured by a Malvern Zetasizer Nano (Malvern Instrument Ltd., UK) with a laser wavelength of 633 nm in 10 mM sodium phosphate buffer at pH 7.4. For the calculation of zeta potential, data processing was done by using Smoluchowski model^40^. For TEM analysis, nanoparticles were loaded onto glow-discharged 200 mesh copper grids (Formvar/carbon grids, Ted Pella) for 20 sec, blot dried, and then stained three times with uranyl formate and dried. TEM imaging was performed using a Tecnai G2 Spirit BioTWIN (FEI) operating at 120 kV acceleration. Images were obtained on a TemCam-F416(R) (TVIPS) CMOS camera. To estimate the size distribution of nanoparticle-corona complexes, the size of at least 150 particles was measured by Fiji/ImageJ. Nanoparticle sizes were determined in aqueous conditions by dynamic light scattering (DLS).

### Quantification of proteins by BCA assay

Proteins on nanoparticles were quantified by using a Pierce BCA Protein Assay Kit (Thermo Scientific). In this protocol, the proteins were quantified without stripping from nanoparticles. The pellet of washed nanoparticle-corona complexes was resuspended in 50 μl PBS. 400 μl of working reagent (copper solution) was added to the samples and standard solutions and then incubated for 30 min at 37 °C. The samples were centrifuged at 20000 g for 30. 200 μl of each supernatant was transferred into a 96 well plate and the absorbance at 562 nm was measured using a Varioscan plate reader (Thermo scientific). In order to evaluate the degree of nanoparticle interference, two control samples were performed: pristine nanoparticles in PBS and in BSA standard solutions ^41^.

### Eluting corona proteins from NPs for electrophoresis and LC-MS/MS analysis

Concentrated 5X Lane Marker reducing sample buffer (SB, Thermo scientific, 0.3 M Tris, 5 % SDS, 50 % Glycerol, and 100 mM DTT) was added to nanoparticle-corona complexes to recover proteins from the NPs. The samples were heated at 95 °C for 5 min to denature and strip off proteins from NPs. DTT as a reducing agent in the sample buffer can cleave the S-S bond in the Sulpho-SASD structure, which helps to cleave the clicked proteins from each other before running them on a SDS-PAGE gel. Then, the samples were centrifuged at 20000 g, 20 min.

### Electrophoresis analysis

For electrophoresis analysis, the recovered proteins were diluted with PBS to adjust the sample buffer and 40 μl of the proteins were separated on a 12 % SDS-polyacrylamide gel (Bolt™ 4-12% Bis-Tris Plus Gels, 10-well) at the constant voltage of 160 V. A PageRuler Unstained Protein Ladder (Thermo scientific) as the molecular weight standard (10-200 kDa) was also run on the gels. For CY5-labeled samples, the gels were first imaged by an Amersham Typhoon NIR laser scanner. Then, the protein bands were detected by Imperial Protein stain (Coomassie brilliant blue, Thermoscientific). The stained gels were scanned on a Bio-Rad gel documentation system. Protein quantification was performed using the plot profile tool in Fiji/ImageJ (ref). The staining intensity and run-length were normalized based on the maximum values.

### LC-MS/MS analysis

The eluted corona proteins from nanoparticles were first precipitated by using the ProteoExtract® Protein Precipitation Kit (Merck, Germany) as described in the manufacturer protocol. The proteins were dissolved in 8 M Urea, 100 mM ammonium bicarbonate with 10 mM DTT. After 30 min adding iodoacetamide to a final concentration of 35 mM alkylated the samples. The alkylation was quenched after 30 min by adding DTT to a final concentration of 35 mM. Subsequently, the samples were diluted 5 times and digested with trypsin 1/50 (w/w) in 16 hr at 37 °C. Tryptic peptides were micropurified using Empore™ SPE C18 Disks packed in 10 μl pipette tips. LC-MS/MS was performed using an EASY-nLC 1000 system (Thermo Scientific) connected to a QExactive+ Mass Spectrometer (Thermo Scientific). Peptides were trapped on a 2 cm ReproSil-Pur C18-AQ column (100 μm inner diameter, 3 μm resin; Dr. Maisch GmbH, Ammerbuch-Entringen, Germany). The peptides were separated on a 15-cm analytical column (75 μm inner diameter) packed in-house in a pulled emitter with ReproSil-Pur C18-AQ 3 μm resin. Peptides were eluted using a flow rate of 250 nl/min and a 20 min gradient from 5% to 35% phase B (0.1% formic acid and 100% acetonitrile). The collected MS files were converted to Mascot generic format (MGF) using Proteome Discoverer (Thermo Scientific). The data were searched against the bovine proteome (uniprot.org). Database search was conducted on a local mascot search engine. The following settings were used: MS error tolerance of 10 ppm, MS/MS error tolerance of 0.1 Da, trypsin as protease, oxidation of Met as a variable modification and carbamidomethyl as a fixed modification.

#### Uni- and multivariate statistical analysis

To test the statistical significance of differences, ANOVA analysis of the data was performed using GraphPad Prism version 8.00 for Windows, GraphPad Software, La Jolla California USA, www.graphpad.com”.

For cluster analysis, the copy number of corona proteins per nanoparticle was used. This approach allows a comparison of different samples without bias for large-sized proteins or the total protein input. Using this method, a protein with a higher copy number in HC+SC than in the four control HC samples is not necessarily considered an SC protein because a higher copy number could be acquired by coincidence. Therefore, the proteins classified as SC are restricted to proteins that had a consistently lower (or zero) copy number in all control samples. Clustered heatmaps were created on square root-transformed and scaled datasets using the packages gplots (ver. 3.0.1.1) and dendextend (ver. 1.9.0) in the R environment (ver. 3.5.1). For unsupervised hierarchical clustering, the distance matrix was calculated using Ward’s minimum variance algorithm with the Euclidean metric.

For each corona protein identified, protein parameters such as the grand average of hydropathy (GRAVY) scores, instability index, and isoelectric point (PI) of the proteins were extracted using ProtParam, a tool available in the SIB ExPASY Bioinformatic Resources Portal ^42^. Principal component analysis (PCA) was performed on scaled datasets using the FactoMineR (ver. 1.41)^43^ and factoextra (ver. 1.0.5) packages for R to explore in a multivariate manner characteristic features of protein parameters among the corona proteins.

### Quantification of individual proteins on nanoparticles

In order to calculate the copy number of individual protein in corona proteins on nanoparticles, three types of measured data were employed:1) emPAI values obtained by LC-MS/MS analysis, 2) quantified total protein mass by BCA assay, and 3) quantified nanoparticles number by reading the fluorescence of nanoparticle. The copy number of proteins per nanoparticles was calculated by using the following expressions ^44^:

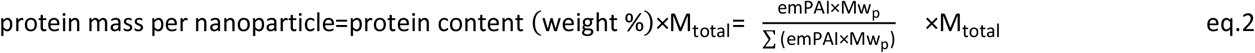

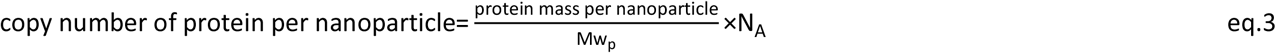

where protein content (weight %) is the contribution of each protein to the total adsorbed mass, Mw_p_ is the calculated molecular weight of the protein, M_total_ is the overall mass of corona proteins per nanoparticles measured by employing BCA assay and fluorescence of nanoparticles. N_A_ is the Avogadro constant (6.023 × 10^23^).

### Estimation of coverage of nanoparticles by corona proteins

In order to estimate the surface coverage of nanoparticles by corona proteins, the Protein Data Bank (PDB) files of the proteins that are available were extracted from PDB:http://www.rcsb.org. Then, the structure of proteins was analyzed by PyMOL and the minimum and maximum cross-section area of proteins were calculated. For the proteins without PDB files, by assuming the simplest shape, sphere, and this partial specific volume (ν=0.73 cm^3^/g), the volume occupied by a protein of mass M in Dalton and its radius were calculated as follows ^45^:

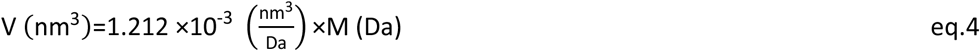

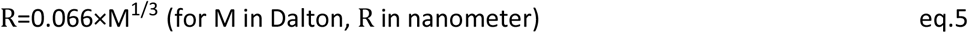

where V and R are the volume and radius of protein.

Since some proteins have quaternary structure and /or are in the lipoprotein structure and it is not clear if the protein prefers to adsorb onto nanoparticle in their natural or denatured structure, it is not possible to calculate the exact coverage of nanoparticles by proteins. On the other hand, proteins can bind to nanoparticles from their maximum and minimum cross-section area. Assuming these limitations for the most abundant proteins, different coverage values were calculated and a range for each calculation was reported.

### Surface plasmon resonance (SPR)

SPR measurements were made on a Biacore 3000 (Biacore AB Sweden). Gold SPR chips from SIA kit Au were cleaned with ultrasonication in acetone, ethanol, and DI water (10 min each), followed by 30 min of UV/ozone, before sputter-deposition of 4 nm Ti followed by 20 nm SiO_2_. The SiO_2_-coated chips were cleaned with ultrasonication in acetone, ethanol, and DI water (10 min each), followed by 30 minutes of UV/ozone a maximum of 1 day before use. All injections were at a rate of 5 μl/min of 100 μl. First, 0.25 mg/ml filtered PLL-g-PEG was injected with 10 mM HEPES (pH 7.4) as a running buffer. Next, 0.25 mg/ml 70 nm SiO_2_ NPs were injected with 10 mM NaCl as a running buffer. The running buffer was changed to 10 mM HEPES containing 100 mM NaCl (pH 7.4) (NaCl-HEPES) for all protein injections. For FBS coated NPs, 1 % FBS in NaCl-HEPES were injected prior to APO H. APO H was injected sequentially, rinsing between each injection, in concentrations 1, 15, 50, 150 μg/ml in NaCl-HEPES.

### SPR data analysis

Linear drift corrections were applied if necessary, using an average of the background drift before and after injection. The two-dimensional fits were made on the MATLAB 2012a platform (Mathworks) using the fitting tool EVILFIT version 3 software ^46,47^ to determine the distribution of binding kinetics. The following input values were used fitting the binding curves: Injection start time: Concentrations: 20 nM, 100 nM, 300 nM, 1000nM and 3000 nM, Start injection: t=0 s, End injection: t=800 s, Fit binding phase from: t=2s, Fit binding phase to: t=798 s, Fit dissociation phase from: t=1400 s, Fit dissociation phase to: 2400 s.

The operator-set boundaries for the distributions were uniformly set to limit K_D_ values in the interval from 10^−9^ to 10^−3^ M, and K_d_ values in the interval from 10^−5^ to 10^0^ s^−1^.

The distribution P (k_a_, K_A_) is calculated using the discretization of the equation:

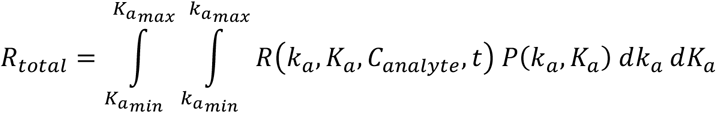

in a logarithmic grid of (k_a_,i, K_a_,i) values with 21 grid points distributed on each axis. This was done through a global fit to association and dissociation traces at the before mentioned analyte concentrations. Tikhonov regularization was used as described by Zhao et al^48^ at a confidence level of P = 0.95 to determine the most parsimonious distribution that is consistent with the data, showing only features that are essential to fit the data.

### Cell culture

We used THP-1 monocyte cells (a human acute monocyte leukemia cell line) obtained from the German Collection of Microorganisms and Cell Cultures (DSMZ, ACC 16) and human cerebral microvessel endothelial hCMEC/D3 cells. Both cell types were grown in RPMI 1640 medium supplemented with 10 % FBS and 1 % penicillin-streptomycin in a humidified 5 % CO_2_ atmosphere at 37 °C. THP-1 cells were differentiated into macrophages (hereafter “THP-1 macrophages”) by incubating with 5 ng/ml PMA for 48 h. The differentiated phenotype was visually inspected under an optical microscope, and then the cells were washed two times with PBS to remove PMA followed by an additional 24 h incubation in the medium without PMA^49^ prior to cell experiments described below.

### Cell association of nanoparticle-corona complexes

Cell association was assessed by a NovoCyte flow cytometer and a confocal laser scanning microscope (CLSM, Zeiss LSM 700.). For flow cytometry, the 0.5 ml of hCMEC/D3 cells or THP-1 macrophages at the density of 5×10^5^ cells/ml were seeded in 24-well plates and exposed to the nanoparticle-corona complexes for 4 h in the RPMI media containing either 5 mg/ml BSA or 10 % FBS. Then, the plates were washed three times with PBS to remove free nanoparticles. The cells were then fixed by 4 % paraformaldehyde for 15 min. The cells were washed 3 times with PBS and resuspended in 200 μl PBS. In the flow cytometry analysis, at least 1000 cells were counted. The fluorescence data are presented as median and calculated as the ratio of the median fluorescence intensity of the samples and the pristine nanoparticles in 5 mg/ml BSA.

For the confocal analysis, first, the glass coverslips were coated with 50 μg/ml collagen type I. For better collagen coating, the coverslips were first coated with poly-d-lysine (PDL) and then with collagen. Then, 2.5×10^5^ THP-1 or hCMEC/D3 cells were seeded onto collagen pre-coated glass coverslips. THP-1 cells were incubated for 48 h with PMA for differentiation followed by 1 day in RPMI without PMA. hCMEC/D3 were incubated for 24 h for attachment. After the differentiation of THP-1 cells and the attachment of hCMEC/D3 cells on coverslips, the cells were exposed to the nanoparticle-corona complexes for 4 h. Then, the cells were washed and fixed with 4 % paraformaldehyde. Cell nuclei were stained with 10 μg/ml Hoechst 33342 (excitation:448 nm; emission:430-480 nm). The actin filaments were stained with 1 μg/ml Phalloidin–Tetramethylrhodamine B isothiocyanate (excitation:540 nm; emission:570-573 nm)

## Acknowledgment

This work acknowledges funding from the FNU project DFF-4181-00473 (The role of soft interactions in complex media) and the Danish National Research Foundation center grant CellPAT (DNRF135).

## Authors contributions

D.S.S., T.V-J., and J.J.E. supervised the project. H.M-B. and D.S.S. conceived and designed the experiments and wrote the manuscript. Y.H performed the statistical and principal component analysis of the mass data and assisted with the preparation of the manuscript. H.M-B., C.M.Z., and H.E. performed the experiments and analysed the data. K.J-M. analysed the SPR data. C.S. performed mass spectroscopy analysis and protein identification. All authors discussed the results and commented on the manuscript.

## References

1. Digiacomo, L., Giulimondi, F., Mahmoudi, M. & Caracciolo, G. Effect of molecular crowding on the biological identity of liposomes: an overlooked factor at the bio-nano interface. Nanoscale Advances (2019).

2. Wang, H. et al. The nature of a hard protein corona forming on quantum dots exposed to human blood serum. Small 12, 5836–5844 (2016).

3. Hayashi, Y. et al. Nanosilver pathophysiology in earthworms: transcriptional profiling of secretory proteins and the implication for the protein corona. Nanotoxicology 10, 303–311 (2016).

4. Monopoli, M. P., Åberg, C., Salvati, A. & Dawson, K. A. Biomolecular coronas provide the biological identity of nanosized materials. Nat Nanotechnol 7, 779–786 (2012).

5. Berardi, A. & Baldelli Bombelli, F. Oral delivery of nanoparticles-let’s not forget about the protein corona. Expert Opin Drug Del 16, 563–566 (2019).

6. Weiss, A. C. et al. Link between Low-Fouling and Stealth: A Whole Blood Biomolecular Corona and Cellular Association Analysis on Nanoengineered Particles. ACS Nano 13, 4980–4991 (2019).

7. Francia, V. et al. Corona Composition Can Affect the Mechanisms Cells Use to Internalize Nanoparticles. ACS Nano 13, 11107–11121 (2019).

8. Giulimondi, F. et al. Interplay of protein corona and immune cells controls blood residency of liposomes. Nat Commun 10, 1–11 (2019).

9. Vu, V. P. et al. Immunoglobulin deposition on biomolecule corona determines complement opsonization efficiency of preclinical and clinical nanoparticles. Nat Nanotechnol 14, 260 (2019).

10. Capjak, I., Goreta, S. S., Jurasin, D. D. & Vrcek, I. V. How protein coronas determine the fate of engineered nanoparticles in biological environment. Arh Hig Rada Toksikol 68, 245–253 (2017).

11. Cedervall, T. et al. Understanding the nanoparticle-protein corona using methods to quantify exchange rates and affinities of proteins for nanoparticles. P Natl A Sci 104, 2050–2055 (2007).

12. Patel, H. Serum opsonins and liposomes: their interaction and opsonophagocytosis. Crit Rev Ther Drug 9, 39–90 (1992).

13. Micluas, T. et al. Dynamic protein coronas revealed as a modulator of silver nanoparticle sulphidation in vitro. Nat Commun 7, 11770 (2016).

14. Fleischer, C. C. & Payne, C. K. Secondary structure of corona proteins determines the cell surface receptors used by nanoparticles. J Phys Chem B 118, 14017–14026 (2014).

15. Wang, J. et al. Soft interactions at nanoparticles alter protein function and conformation in a size dependent manner. Nano Lett 11, 4985–4991 (2011).

16. Miclaus, T., Bochenkov, V. E., Ogaki, R., Howard, K. A. & Sutherland, D. S. Spatial mapping and quantification of soft and hard protein coronas at silver nanocubes. Nano Lett 14, 2086–2093 (2014).

17. Weiss, A. C. et al. In Situ Characterization of Protein Corona Formation on Silica Microparticles Using Confocal Laser Scanning Microscopy Combined with Microfluidics. ACS Appl Mater Inter 11, 2459–2469 (2019).

18. Lu, X. et al. Tailoring the component of protein corona via simple chemistry. Nat Commun 10, 1–14 (2019).

19. Yan, Y. et al. Differential roles of the protein corona in the cellular uptake of nanoporous polymer particles by monocyte and macrophage cell lines. ACS Nano 7, 10960–10970 (2013).

20. Vroman, L. & Adams, A. L. Identification of rapid changes at plasma-solid interfaces. J Biomed Mater Res 3, 43–67 (1969).

21. Walczyk, D., Bombelli, F. B., Monopoli, M. P., Lynch, I. & Dawson, K. A. What the cell “sees” in bionanoscience. J Am Chem Soc 132, 5761–5768 (2010).

22. Liu, X. et al. Rapid conjugation of nanoparticles, proteins and siRNAs to microbubbles by strain-promoted click chemistry for ultrasound imaging and drug delivery. Polym Chem 10, 705–717 (2019).

23. DeForest, C. A. & Anseth, K. S. Cytocompatible click-based hydrogels with dynamically tunable properties through orthogonal photoconjugation and photocleavage reactions. Nat Chem 3, 925 (2011).

24. Ishizuka, T., Liu, H. S., Ito, K. & Xu, Y. Fluorescence imaging of chromosomal DNA using click chemistry. Sci Rep 6, 33217 (2016).

25. Schreiber, C. L. & Smith, B. D. Molecular conjugation using non-covalent click chemistry. Nat Rev Chem 3, 393–400 (2019).

26. Wang, S. et al. Regulation of Ca 2 Signaling for Drug-Resistant Breast Cancer Therapy with Mesoporous Silica Nanocapsule Encapsulated Doxorubicin/siRNA Cocktail. ACS Nano 13, 274–283 (2018).

27. Li, L. et al. Polystyrene nanoparticles reduced ROS and inhibited ferroptosis by triggering lysosome stress and TFEB nucleus translocation in a size-dependent manner. Nano Lett 19, 7781–7792 (2019).

28. Llopis-Lorente, A. et al. Enzyme-Powered Gated Mesoporous Silica Nanomotors for On-Command Intracellular Payload Delivery. ACS Nano 13, 12171–12183 (2019).

29. Docter, D. et al. Quantitative profiling of the protein coronas that form around nanoparticles. Nat Protoc 9, 2030–2044 (2014).

30. Carril, M. et al. In situ detection of the protein corona in complex environments. Nat Commun 8, 1542 (2017).

31. Zeuthen, C. M., Shahrokhtash, A. & Sutherland, D. S. Nanoparticle adsorption on antifouling polymer brushes. Langmuir 35, 14879–14889 (2019).

32. Tenzer, S. et al. Rapid formation of plasma protein corona critically affects nanoparticle pathophysiology. Nat Nanotechnol 8, 772–81 (2013).

33. Lunov, O. et al. Differential uptake of functionalized polystyrene nanoparticles by human macrophages and a monocytic cell line. ACS Nano 5, 1657–69 (2011).

34. Platt, N. & Gordon, S. Scavenger receptors: diverse activities and promiscuous binding of polyanionic ligands. Chem Biol 5, R193–R203 (1998).

35. Casals, E., Pfaller, T., Duschl, A., Oostingh, G. J. & Puntes, V. Time evolution of the nanoparticle protein corona. ACS Nano 4, 3623–32 (2010).

36. Pyzik, M. et al. The Neonatal Fc Receptor (FcRn): A Misnomer? Front Immunol 10, 1540 (2019).

37. Urich, E., Lazic, S. E., Molnos, J., Wells, I. & Freskgård, P.-O. Transcriptional profiling of human brain endothelial cells reveals key properties crucial for predictive in vitro blood-brain barrier models. PloS one 7, e38149 (2012).

38. Ritz, S. et al. Protein corona of nanoparticles: distinct proteins regulate the cellular uptake. Biomacromolecules 16, 1311–1321 (2015).

39. Thiagarajan, P., Le, A. & Benedict, C. R. Beta(2)-glycoprotein I promotes the binding of anionic phospholipid vesicles by macrophages. Arterioscler. Thromb. Vasc. Biol. 19, 2807–11 (1999).

40. Hunter, R. J. Zeta potential in colloid science: principles and applications. 2, (Academic press, 2013).

41. Monopoli, M. P., Pitek, A. S., Lynch, I. & Dawson, K. A. Formation and characterization of the nanoparticle-protein corona. Nanomaterial Interfaces in Biology: Methods and Protocols 137–155 (2013).

42. Gasteiger, E. et al. ExPASy: the proteomics server for in-depth protein knowledge and analysis. Nucleic Acids Res 31, 3784–3788 (2003).

43. Lê, S., Josse, J. & Husson, F. FactoMineR: An R package for multivariate analysis. J. Stat. Softw 25, 1–18 (2008).

44. Ishihama, Y. et al. Exponentially modified protein abundance index (emPAI) for estimation of absolute protein amount in proteomics by the number of sequenced peptides per protein. Mol Cell Proteomics 4, 1265–1272 (2005).

45. Erickson, H. P. Size and shape of protein molecules at the nanometer level determined by sedimentation, gel filtration, and electron microscopy. Biological procedures online 11, 32 (2009).

46. Vorup-Jensen, T. in Integrin and Cell Adhesion Molecules 55–71 (Springer, 2011).

47. Jensen, M. R. et al. Structural basis for simvastatin competitive antagonism of complement receptor 3. J. Biol. Chem. 291, 16963–16976 (2016).

48. Zhao, H., Gorshkova, I. I., Fu, G. L. & Schuck, P. A comparison of binding surfaces for SPR biosensing using an antibody-antigen system and affinity distribution analysis. Methods 59, 328–35 (2013).

49. Park, E. et al. Optimized THP-1 differentiation is required for the detection of responses to weak stimuli. Inflamm. Res. 56, 45–50 (2007).

