## supplementary information for "Mapping and identification of soft corona proteins at nanoparticles and their impact on cellular association"

### Supplementary tables

**Supplementary Table S1. Degree of labelling of proteins with DBCO Sulpho-NHS and Sulpho-NHS CY5**

| | Condition | | Degree of labeling (DOL) | |
|---|---|---|---|---|
| | **DBCO (mM)** | **Sulpho CY5 (mM)** | **DBSO** | **CY5** |
| **FBS-D** | 0.2 | 0 | 4.2±0.3 | - |
| | 0.4 | 0 | 5.1±0.2 | - |
| | 0.8 | 0 | 5.6±0.1 | - |
| **FBS-CY5** | 0 | 0.13 | - | 1.1±0.08 |
| **FBS-D-CY5** | 0.4 | 0.13 | 4.9±0.1 | 1.2±0.1 |

**Supplementary Table S1. Calculation of degree of labeling (DOL) of proteins.** The degree of labelling (DOL) of proteins were calculated by equation 1, using the UV-Vis absorbance of samples at 280 nm, 309 nm, and 646 nm for proteins, DBCO, and CY5, respectively.

**Supplementary Table S2. Characterization of nanoparticle-corona complexes**

| | nanoparticle-corona complexes | Zeta potential ± SD (mV) | DLS hydrodynamic diameter ± SD (nm) (PDI) | | nanoparticle-corona complexes | Zeta potential ± SD (mV) | DLS hydrodynamic diameter ± SD (nm) (PDI) |
|---|---|---|---|---|---|---|---|
| SNPs | pristine | -22±3.4 | 81.9±5.1 (0.01) | PsNPs | pristine | -27.3±2.1 | 110±11.2 (0.02) |
| | HC | -20±4.1 | 106±5.7 (0.06) | | HC | -26.2±2.3 | 155±8.3 (0.1) |
| | HC-$N_3$ | -25±2.4 | 128±6.3 (0.12) | | HC-$N_3$ | -25.4±3.1 | 168±14.2 (0.15) |
| | D Ctrl | -23±3.7 | 133±7.5 (0.08) | | D Ctrl | -23.6±2.8 | 179±13.6 (0.19) |
| | $N_3$ Ctrl | -26±3.3 | 121±10.2 (0.07) | | $N_3$ Ctrl | -26.7±4.3 | 175±9.2 (0.2) |
| | HC+SC | -29±2.9 | 152±12.4 (0.17) | | HC+SC | -25.3±4.2 | 191±15.3 (0.18) |

**Supplementary Table S2.** The average size of nanoparticle-corona complexes was determined using DLS and the zeta potential measurement data processing was done by using Smoluchowski model. Zeta potential measurement was done in 10 mM sodium phosphate buffer, pH 7.4, containing 10 mM NaCl. Data shown correspond to mean ± sd. of three independent experiments (n=3).

**Supplementary Table S3.** Kinetic parameters of binding of APO H to HC proteins on SNPs.

| Population | % of total signal | Koff ($s^{-1}$) | Kd (M) |
|---|---|---|---|
| **1** | 88 ± 4.4 | $4.8\times10^{-2} \pm 8.0\times10^{-3}$ | $1.7 \times10^{-5} \pm 3.6\times10^{-6}$ |
| **2** | 9.54 ±3.46 | $8.1\times10^{-5} \pm 4.9\times10^{-6}$ | $7.3\times10^{-8} \pm 6.6\times10^{-9}$ |

**Supplementary Table S3.** Two-dimensional fits were applied to the SPR data to achieve values for Kd and Koff for different populations of APO H binding to the HC on SNPs.

### Supplementary Figure

#### Supplementary Fig. S1.

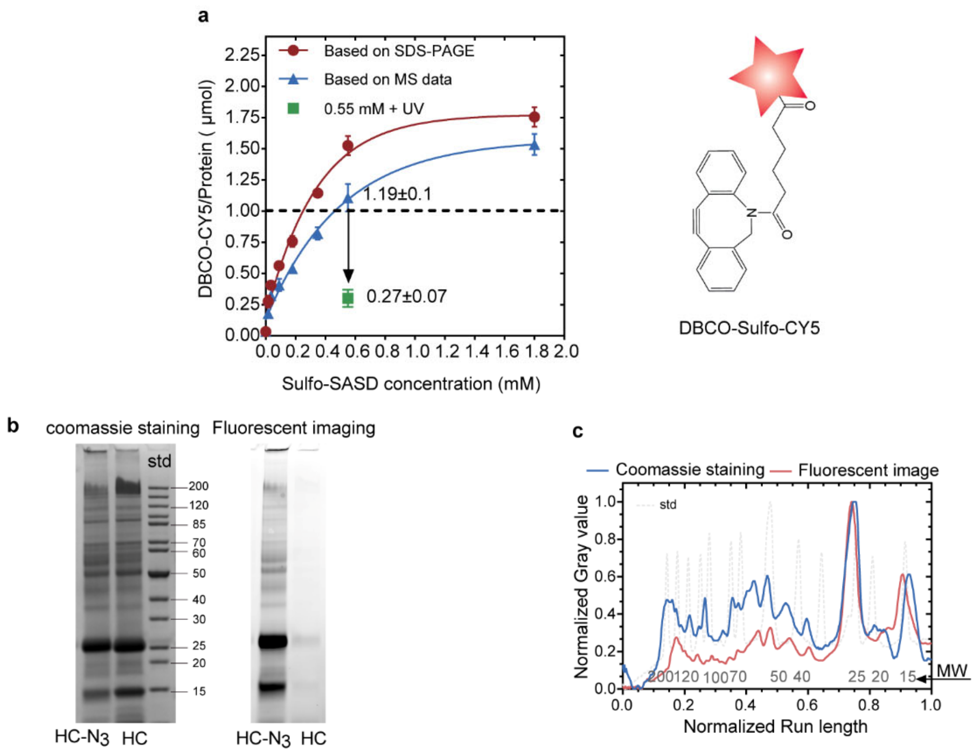


**Supplementary Fig. S1. Optimization of modification of HC proteins on SNPs with Sulpho-SASD.** SNPs-hard corona complexes (0.4 mg/ml) were incubated with different concentrations of Sulpho-SASD (0, 0.018, 0.036, 0.09, 0.18, 0.35, 0.55, 1.8 mM) for 1 h, Sulpho-SASD modifies protein through the reaction of its Sulpho-NHS with primary amines on proteins. To confirm the labelling, the azide-modified particles were incubated with DBCO-Sulpho-CY5 which reacts with the azide groups through a SPAAC click reaction. The labelling efficiency was measured using two methods. In the first method, the amount of protein was calculated by a combination of SDS-PAGE and BCA assay. In the second method, the LC-MS/MS data was used to calculate the protein content on SNPs. Converting N3 group by UV to nitren decreased the click reaction efficiency, which is considered as another control experiment. Sulpho-SASD at 0.6 mM was used for further steps. **b,c,** $N_3$ modification of HC proteins formed on SNPs over 2 h incubation with FBS was characterized by a click reaction between $N_3$ and DBCO-Sulpho-CY5. Fluorescence image of the SDS-PAGE (**b**) and the comparison between the densitometry of coomassie staining and fluorescence image (**c**) show that all the proteins stained with coomassie reacted with DBCO, which confirms the presence of $N_3$ on all HC proteins.

**Supplementary Fig. S2.**

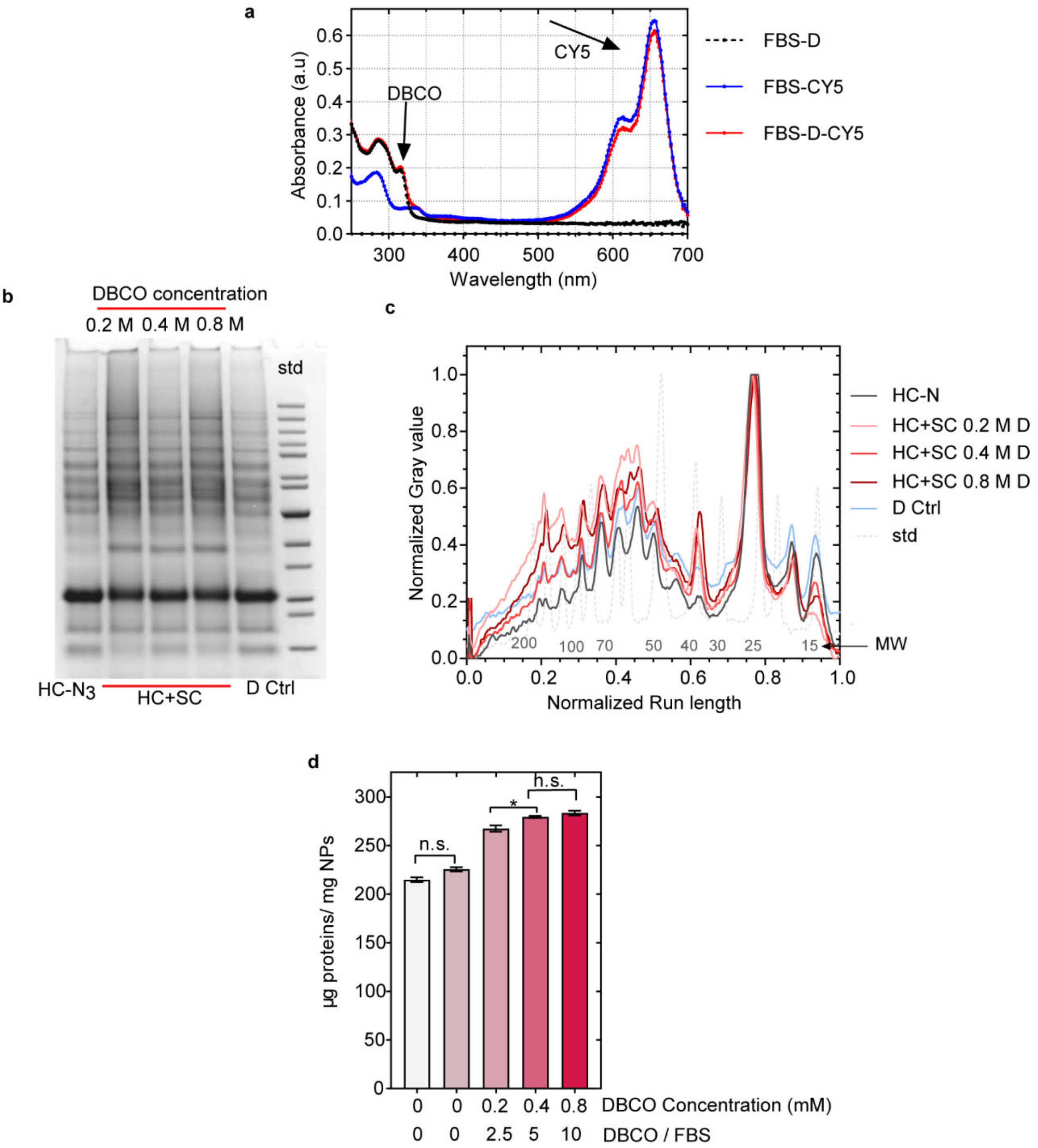


**Supplementary Fig. S2. Optimization of SPAAK "click" reaction. a,** UV-vis spectroscopy analysis of FBS proteins labelled with DBCO or CY5. The degree of labelling (DOL) in the Supplementary Table.S1 show that labelling efficiency does not change when the proteins were labelled with both DBCO and CY5. **b-d,** Different DBCO concentrations (0, 0.2, 0.4, and 0.8 mM equals to 0, 2.5, 5, and 10 of DBCO/FBS) were used for modification of FBS proteins. The corona protein was visualized by a SDS-PAGE gel (**b**), analysed with densitometry analysis (**c**), and quantified by BCA assay (**d**). The SDS-PAGE analysis shows addition of some proteins by the click reactions. The BCA assay shows that increasing DBCO concentration from 0.4 to 0.8 mM did not lead to a significant change in the amount of protein corona. Nomenclature: FBS-D: FBS proteins modified with DBCO, hard corona (HC), hard corona modified with azide (HC-$N_3$), FBS-D added to HC (D Ctrl), FBS added to HC-N3 ($N_3$ Ctrl), FBS-D added to HC-$N_3$ (HC_SC), and FBS modified with both CY5 and DBCO (FBS-D-CY5).

**Supplementary Fig. S3.**

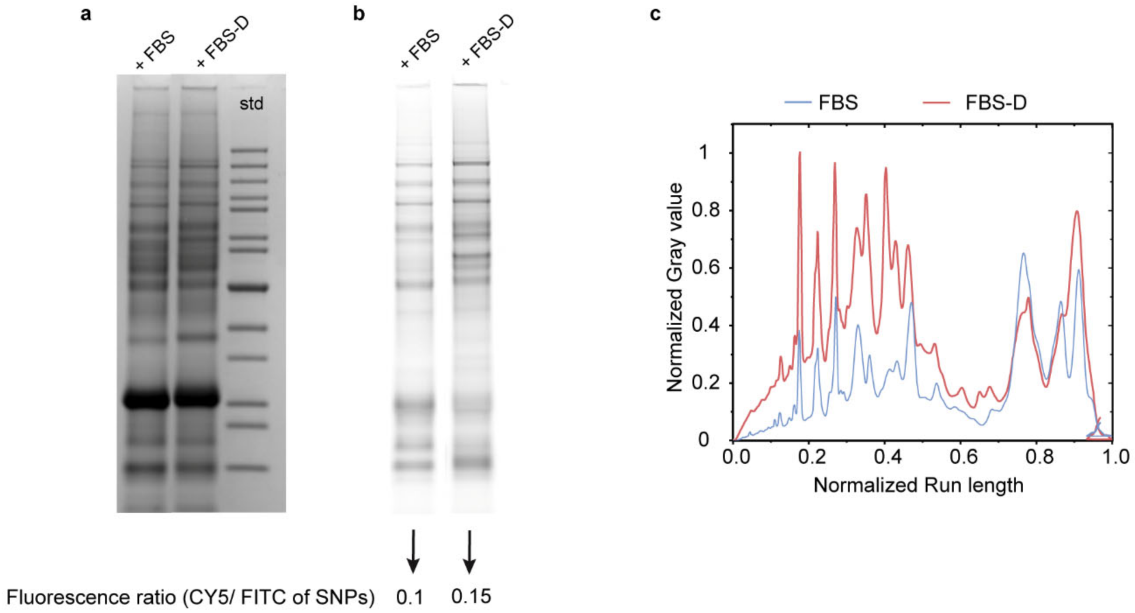


**Supplementary Fig. S3. Addition of fluorescently labelled FBS to SNPs@HC-$N_3$. a-c,** Coomassie staining (**a**), fluorescence image (**b**), and densitometry analysis of the fluorescence image of SDS-PAGE gel of fluorescently labelled FBS and FBS-D proteins added to SNPS@HC-$N_3$. Densitometry analysis clearly shows the addition of more proteins through click chemistry to HC on SNPs. The fluorescence ratio of CY5/FITC of SNPs was also measured in both conditions (showed below the SDS-PAGE image) which confirms the addition of more fluorescently labelled proteins to HC through click chemistry.

**Supplementary Fig. S4.**

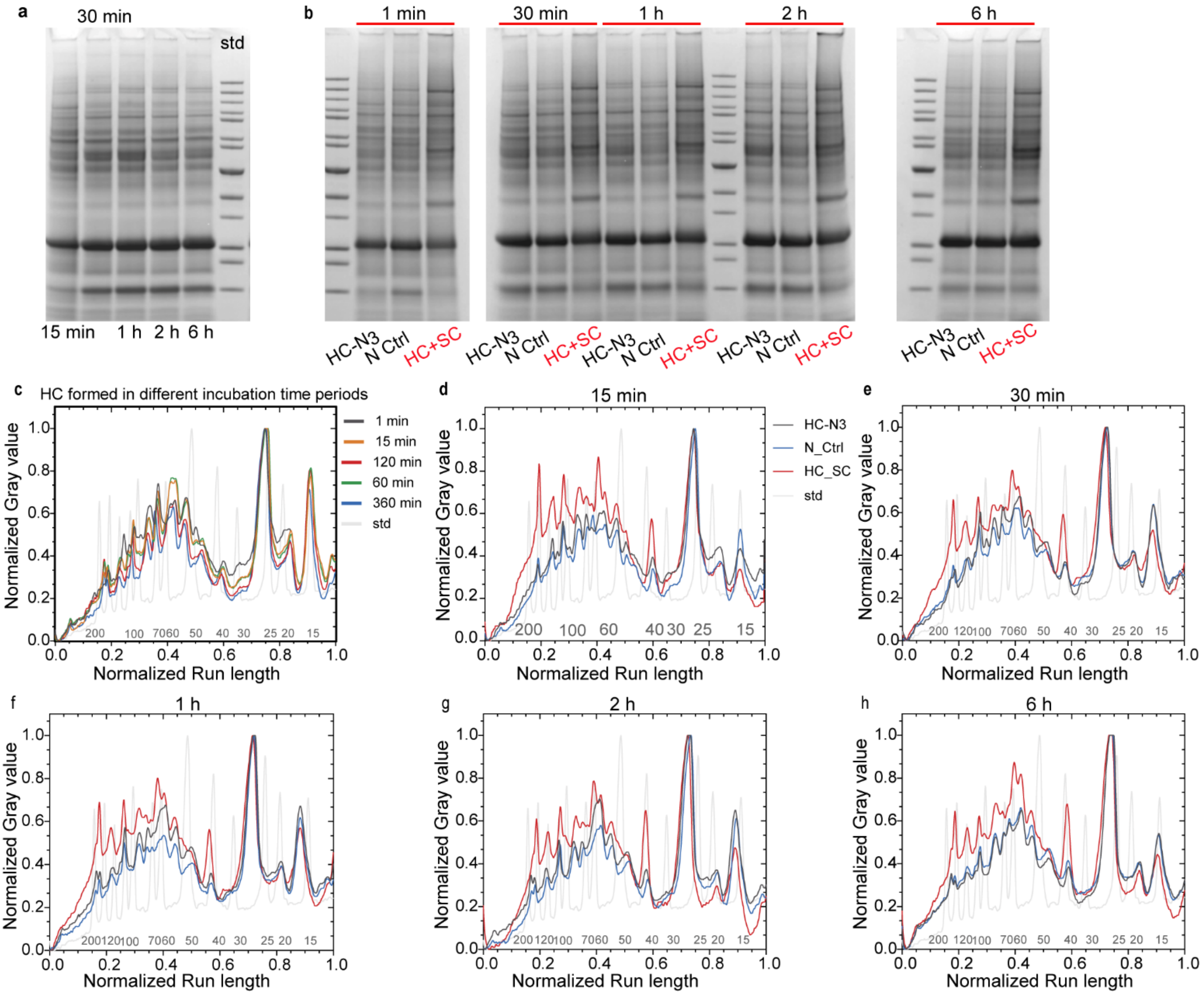


**Supplementary Fig. S4. Capturing weakly interacting proteins on HC proteins which were formed by exposure to FBS for the indicated time periods (15 min, 30 min, 1h, 2h, and 6 h). a,b,** SDS-PAGE of HC formed on SNPs over different exposure time periods (**a**) and HC+SC proteins on SNPs **(b). c**, densitometry analysis of SDS-PAGE gel in (**a**). **d-h,** densitometry analysis of SDS-PAGE gels shown in (**b**).

**Supplementary Fig. S5**

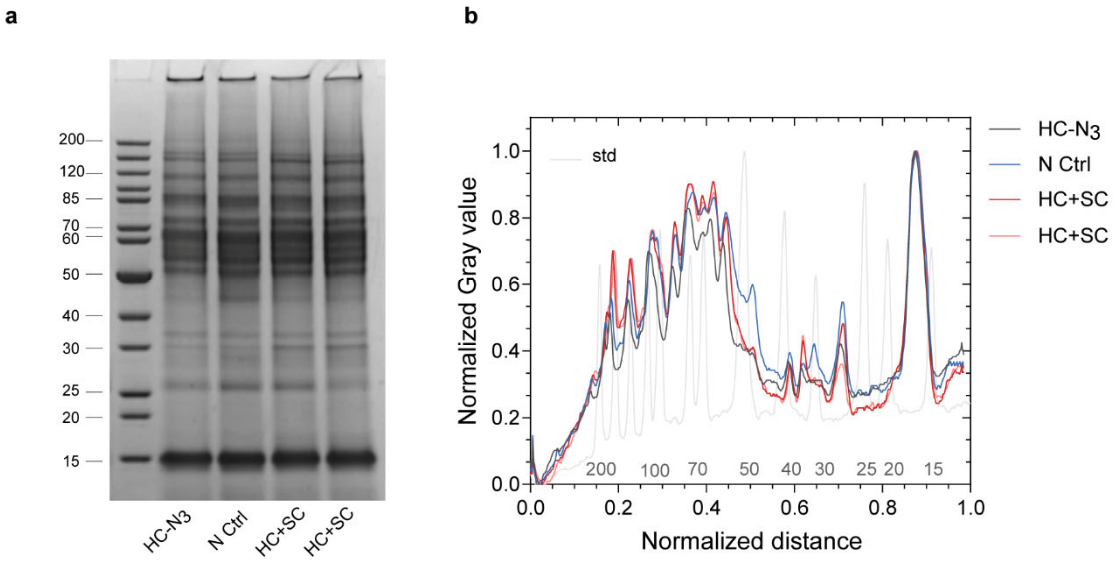


**Supplementary Fig. S5. a,b,** SDS-PAGE image **(a)** and densitometry analysis **(b)** of eluted corona proteins (HC and HC+SC) from PsNPs which were captured through click chemistry reaction.

**Supplementary Fig. S6.**

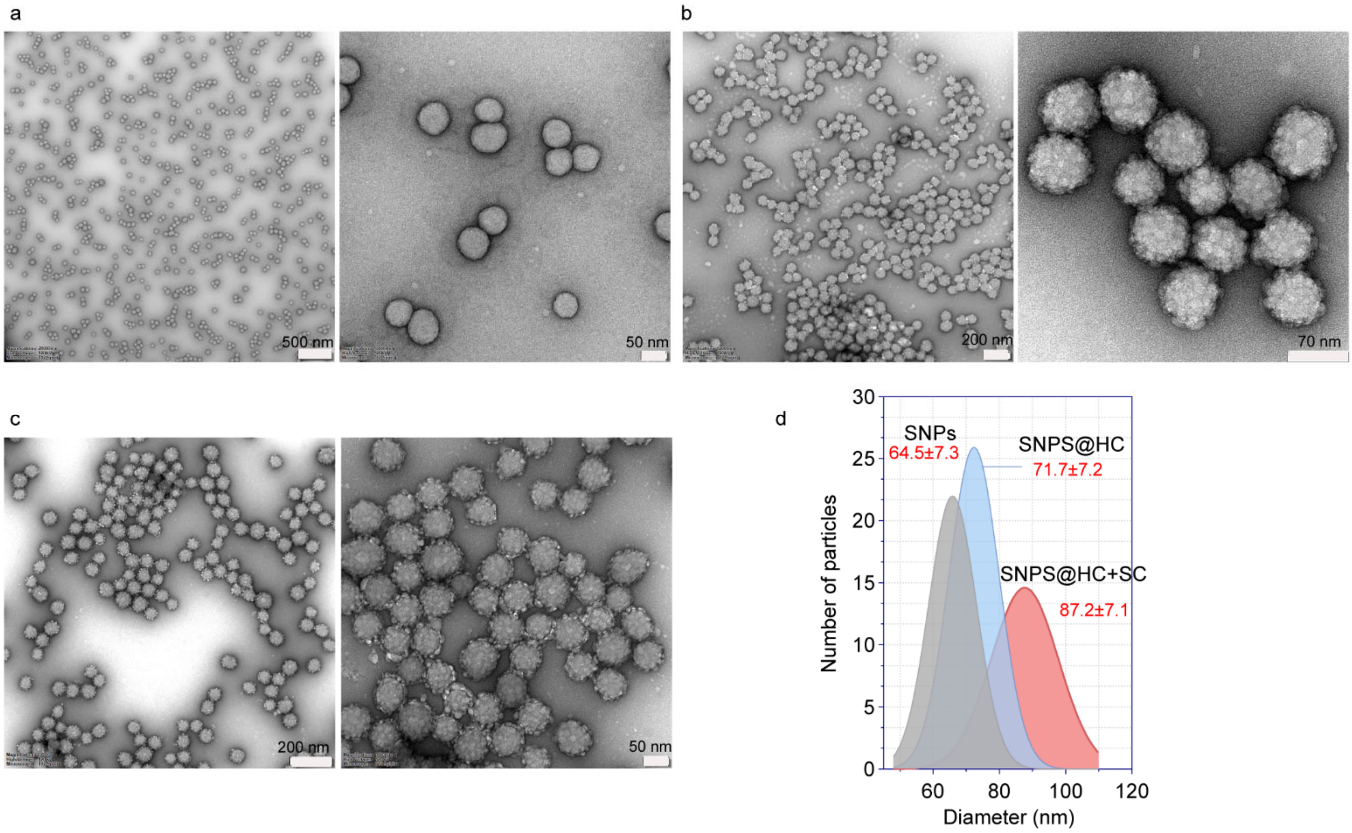


**Supplementary Fig. S6.** TEM analysis of pristine SNPs (**a**), SNPs@HC (**b**), and SNPs@HC+SC (**c**). **d**, The average size of nanoparticles was calculated by measuring the size of at least 150 particles.

**Supplementary Fig. 7.**

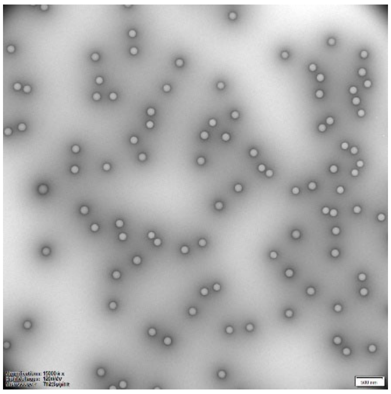

**Supplementary Fig. S7.** TEM analysis of pristine PsNPs. Scale bar, 500 nm.

**Supplementary Fig. S8.**

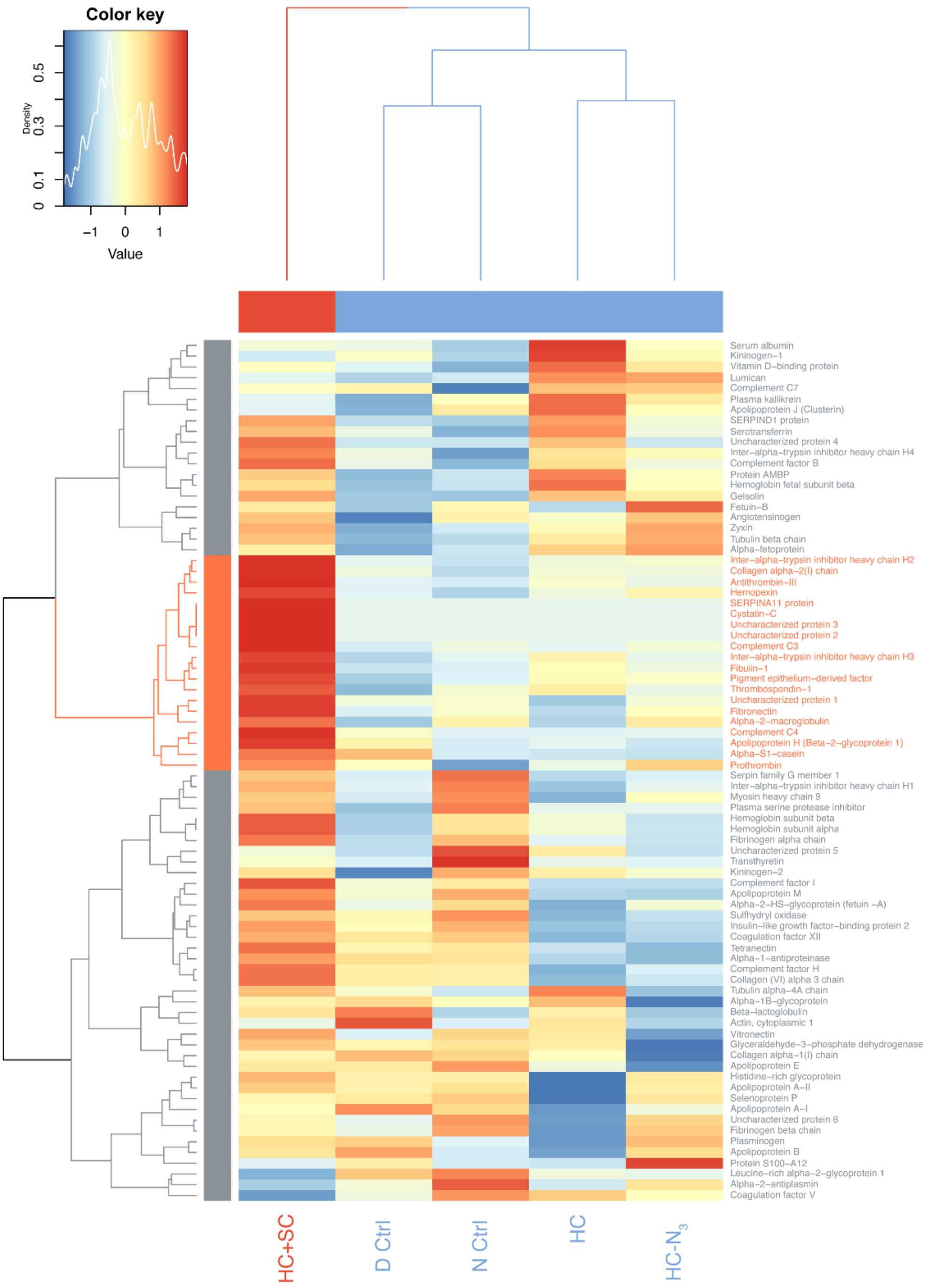


**Supplementary Fig. S8.** A heatmap with two-way unsupervised hierarchical clustering analysis (UHCA) of the relative abundance of corona proteins recovered from SNPs. Each row, a protein; each column, a protein corona sample. The number of proteins per nanoparticle is scaled to derive a z-score representing the relative abundance of each protein between the samples. A colour key along with the z-score distribution is depicted to the top left. Red and blue correspond to the number of proteins higher and lower than the average across all samples, respectively. The row dendrogram reveals a putative SC cluster (coloured in orange) characterized by specific enrichment of the proteins in HC+SC.

**Supplementary Fig. S9.**

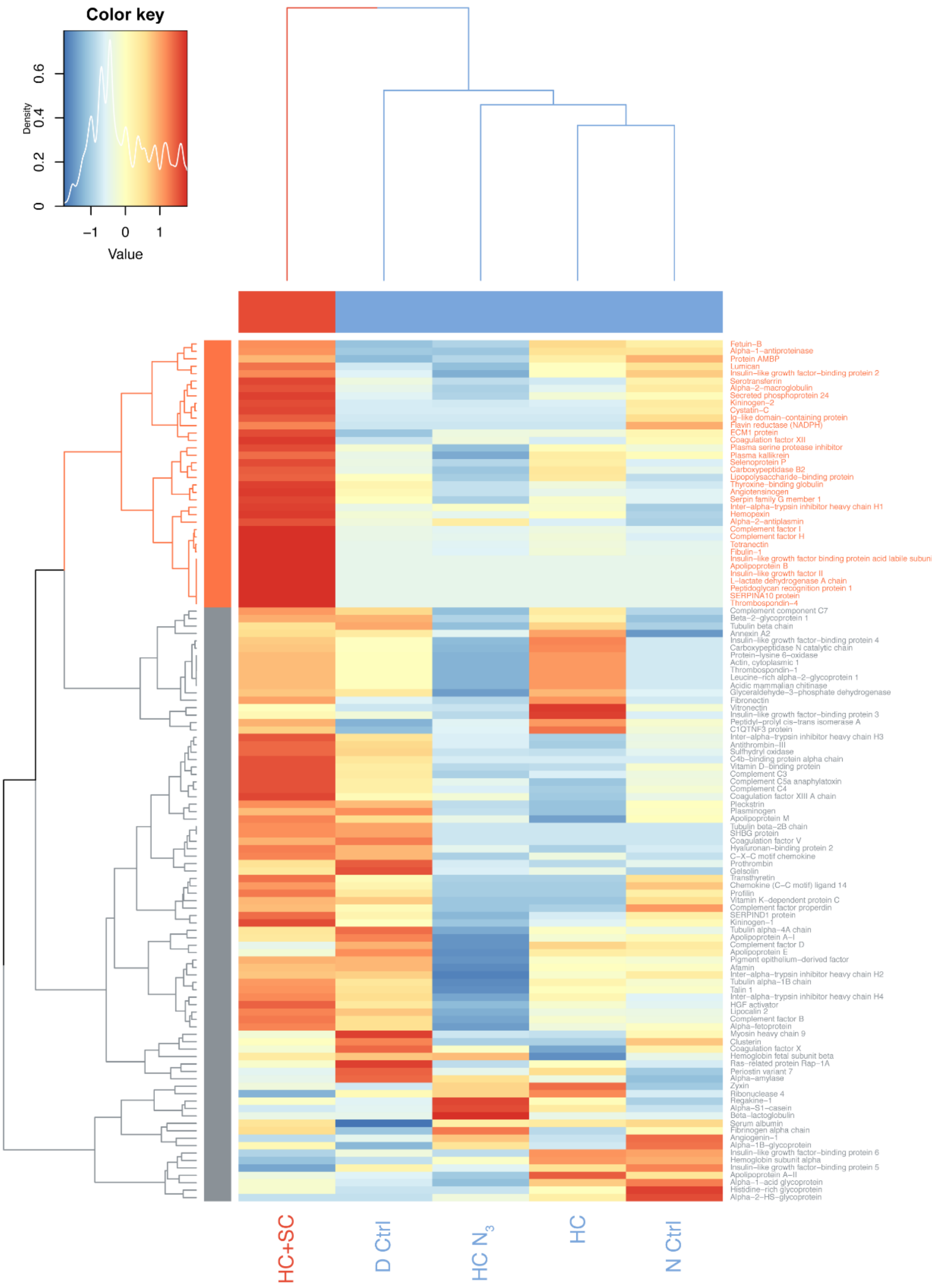


**Supplementary Fig. S9.** A heatmap with two-way unsupervised hierarchical clustering analysis (UHCA) of the relative abundance of corona proteins recovered from SNPs. Each row, a protein; each column, a protein corona sample. The number of proteins per nanoparticle is scaled to derive a z-score representing the relative abundance of each protein between the samples. A colour key along with the z-score distribution is depicted to the top left. Red and blue correspond to the number of proteins higher and lower than the average across all samples, respectively. The row dendrogram reveals a putative SC cluster (coloured in orange) characterized by specific enrichment of the proteins in HC+SC.

**Supplementary Fig. S10.**

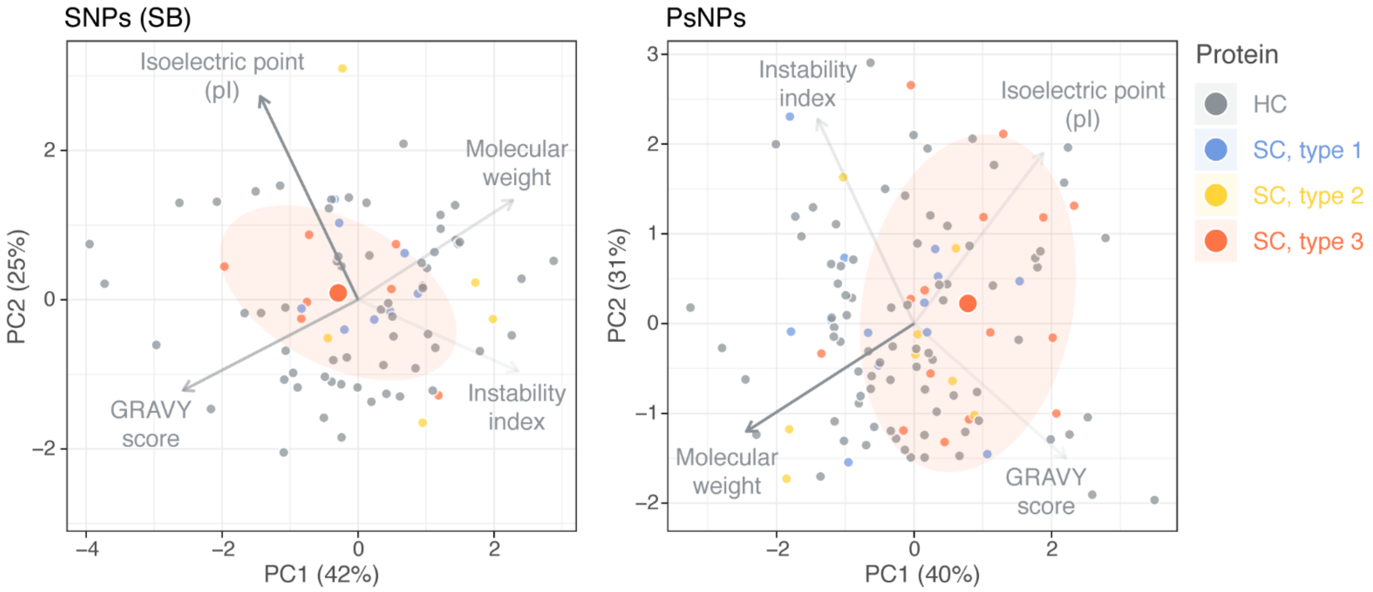


**Supplementary Fig. S10. Parameter analysis of corona proteins eluted from SNPs and PsNPs.**

**Supplementary Fig. S11.**

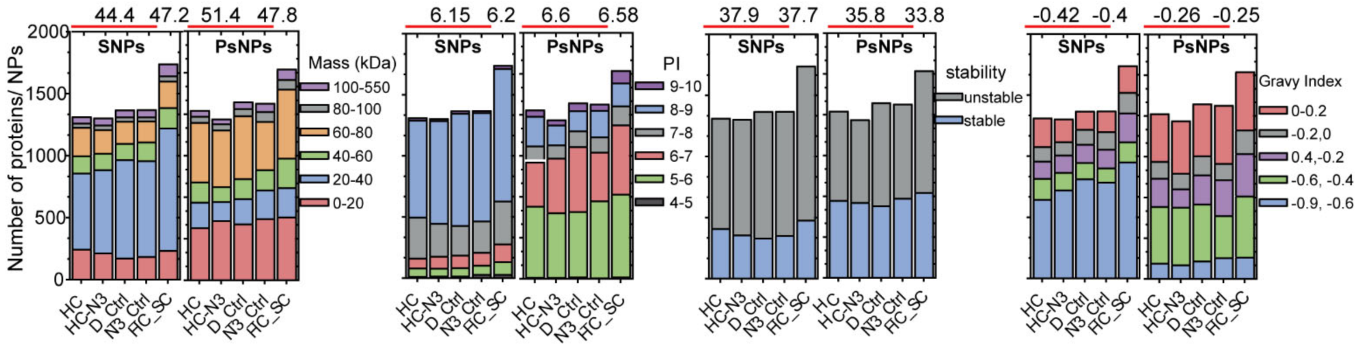


**Supplementary Fig. S11. Parameter analysis of corona proteins eluted from SNPs (LB).** Classification of corona proteins identified in different SNPs and PsNPs samples by LC_MS/MS according to their calculated molecular mass isoelectric point, instability index, and gravy index. The number-weighted protein parameters written above the figures were calculated for HC (average of four control samples) and HC+SC. It should be mentioned that in spite of almost the same negative charge on both SNPs and PsNPs, the classification of corona proteins on SNPs and PsNPs was different, as thoroughly discussed in the previous studies [1,2]. This shows that other than electrostatic interactions between proteins and NPs with opposite charges, there can be electrostatic interactions between the exposed part of a denatured protein with the same charge, and nanoparticles, or other interactions such as hydrophobic interactions [3,4].

**Supplementary Fig. S12.**

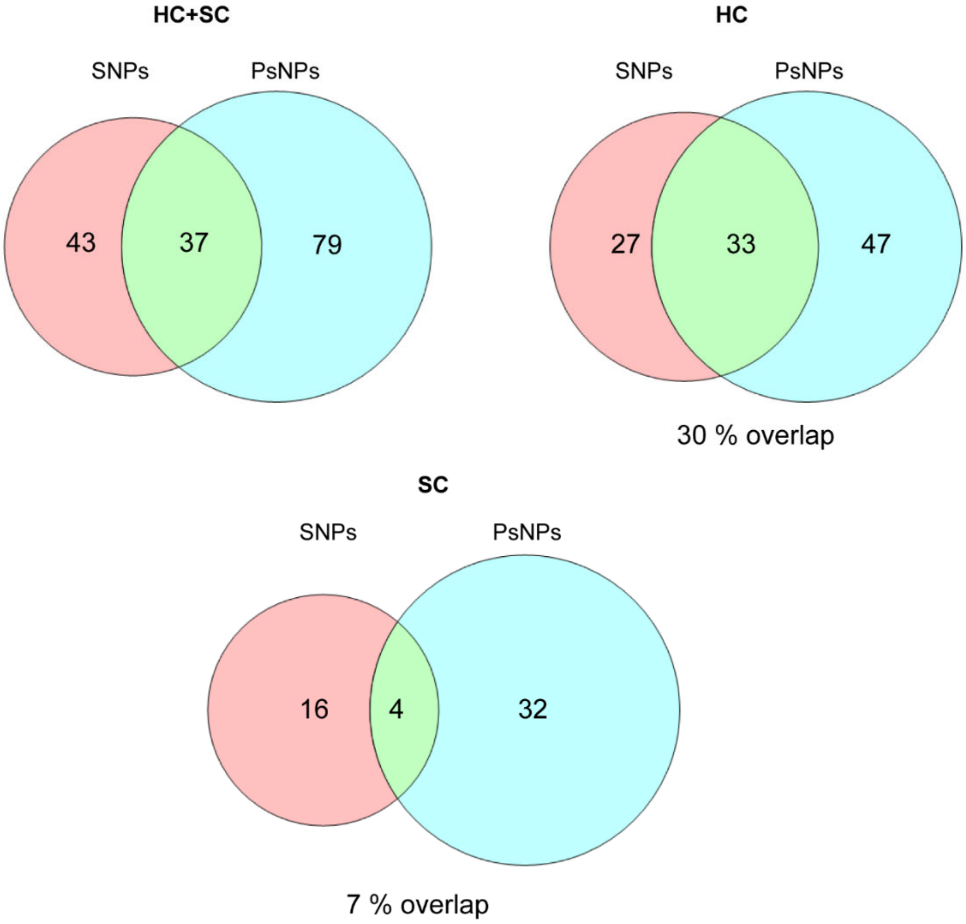


**Supplementary Fig. S12. Venn diagrams depicting the degree of overlap of corona proteins on SNPs and PsNPs.** HC+SC (total corona proteins), HC (hard corona proteins), and SC (soft corona proteins).

**Supplementary Fig. S13**

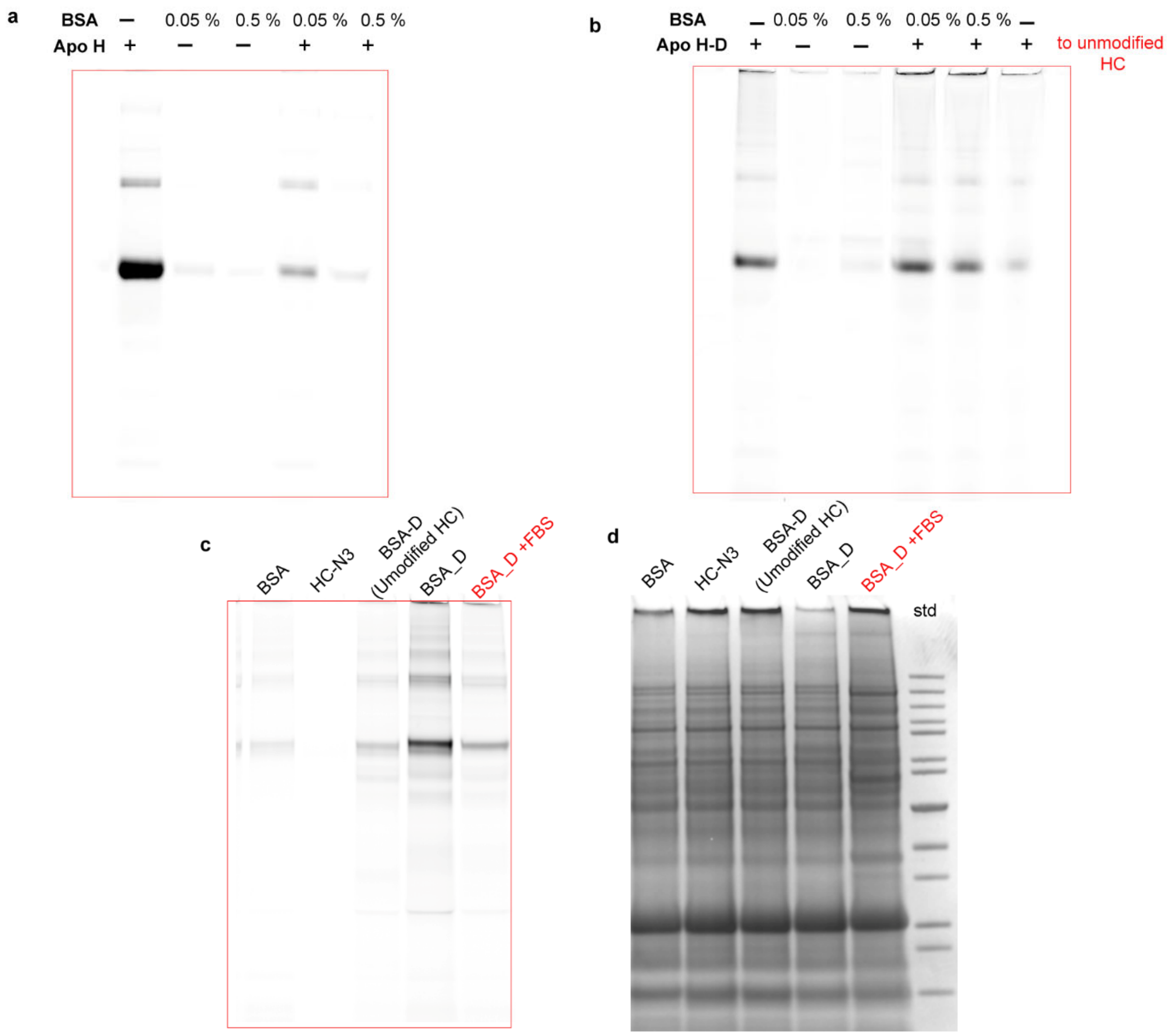


**Supplementary Fig. S13. The complete SDS-PAGE of competition study shown in Fig. 3. a,** Fluorescence image of SDS-PAGE gel of proteins eluted from nanoparticles in the experiment of the addition of fluorescently labeled APO H to SNPs (**a**) and APO H-D to SNPs@HC-$N_3$ (**b**) in the presence of varying concentrations of BSA (0.05 and 0.5 %). **c,d,** fluorescence image (**c**) and coomassie stained image (**d**) of the addition of fluorescently labeled BSA to SNPs@HC-$N_3$ in the presence and absence of FBS proteins.

**Supplementary Fig. S14.**

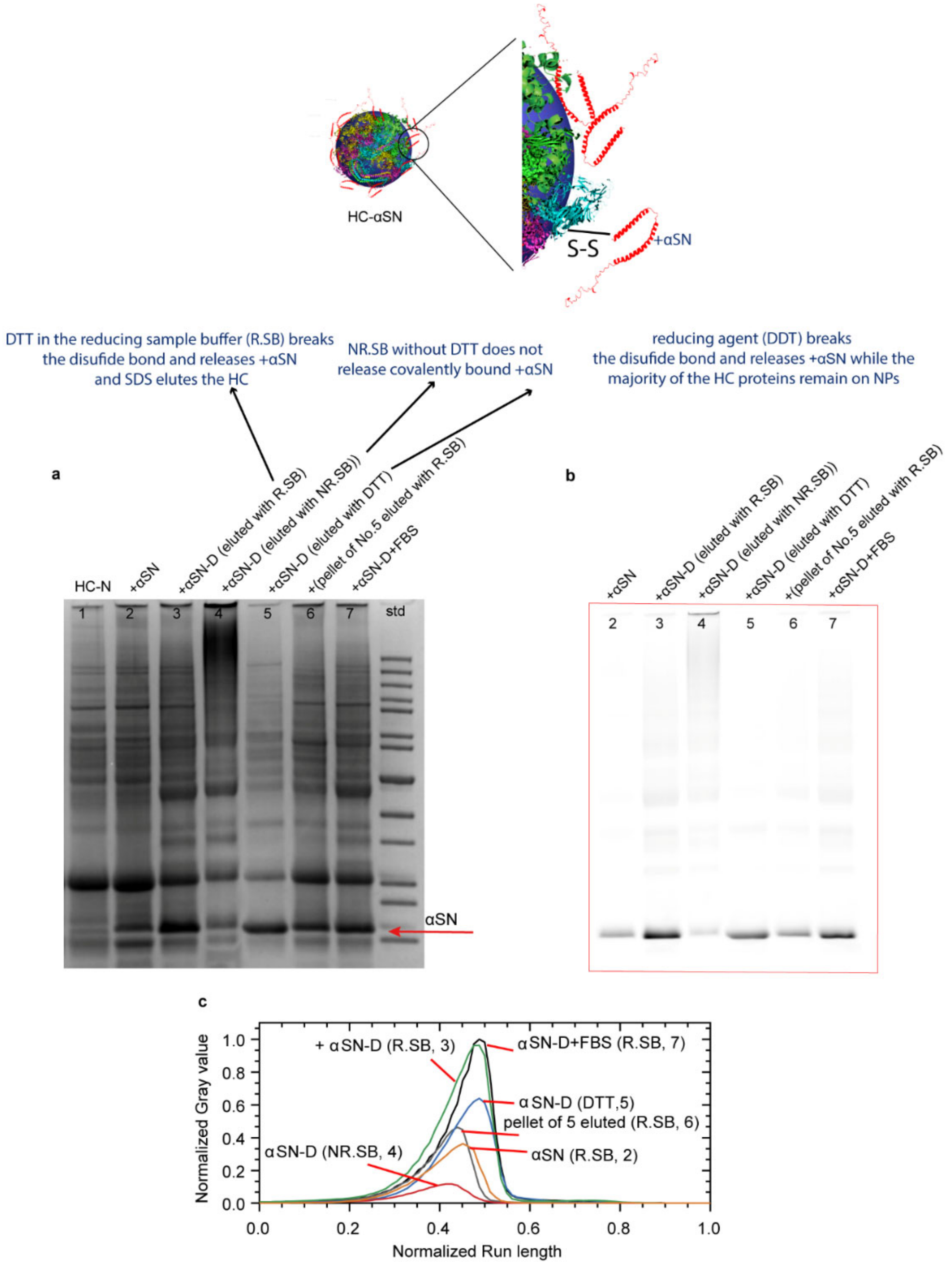


**Supplementary Fig S14. Capturing fluorescently labeled α-Synuclein (αSN) as a disease-related protein on HC on SNPs by click chemistry reaction. The corona proteins were eluted with different elution buffers, reducing sample buffer containing SDS, DTT, and glycerol (R.SB), non-reducing sample buffer containing SDS and glycerol (NR.SB), and DTT alone as a reducing agent. a-c,** coomassie stained (**a**) and fluorescent image (**b**) of a SDS-PAGE gel, and densitometry analysis of the fluorescent image of SDS-PAGE gel of proteins eluted from SNPs. Numbers in the images are as follows: 1) HC-N3 (eluted with R.SB), 2) αSN added to HC-N3 on SNPs (eluted with R.SB), 3) αSN-D added to HC-N3 on SNPs (eluted with R.SB), 4) αSN-D added to HC-N3 on SNPs (eluted with NR.SB), 5) αSN-D added to HC-N3 on SNPs (eluted with DTT alone), 6) proteins on the pellet of No.5 was eluted with R.SB, 7) αSN-D added to HC-N3 on SNPs in the presence of FBS (eluted with R.SB). The results confirm that DTT is necessary to reduce the disulfide bridge in the Sulpho-SASD structure to release the majority of αSN.

**Supplementary Fig. S15.**

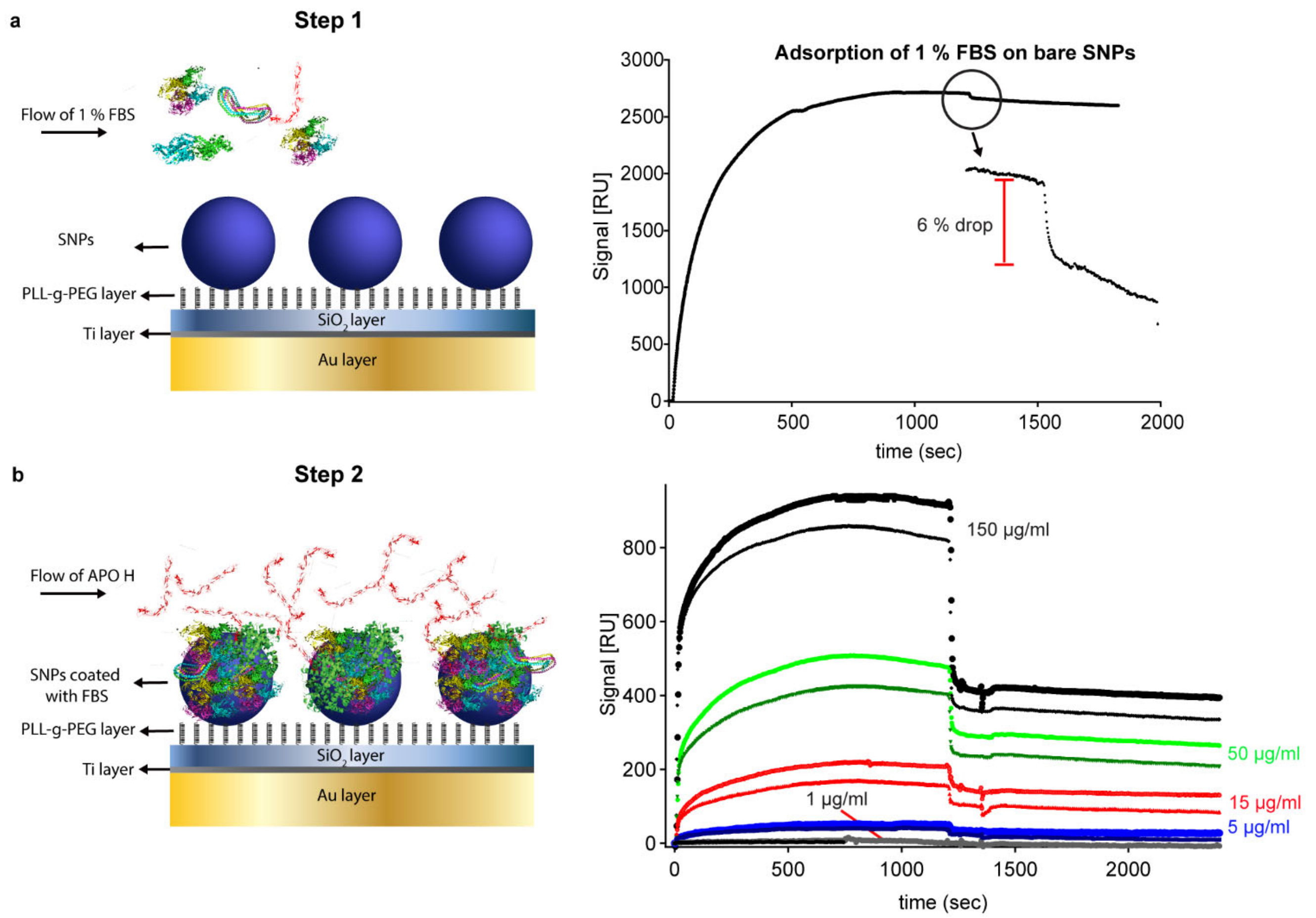


**Supplementary Fig. S15. a,** SPR measurements on SNPs on PLL-g-PEG with injections of 1 % FBS. The protein corona was formed by injecting 1 % FBS onto the immobilized SNPs. The FBS binds to the NPs, and reaches a stable plateau during this injection signifying equilibrium has been reached. The signal only drops 6% when rinsing commences, indicating that the proteins are tightly adsorbed to the NPs. **b,** Injection of 1, 5, 15, 50 and 150 µg/ml APO H following the FBS injection. A rapid drop of up to 60 % of the adsorbed amount of ApoH was seen in 50 µg/ml APO H immediately upon rinsing. This indicates that a fraction of the APO H associates with the NPs as part of the soft corona that is instantly removed by rinsing. A fraction of the APO H stays associated with the NPs even after rinsing, which was also expected based on a gel with HC + APO H incubation (Fig.3).

**Supplementary Fig. S16.**

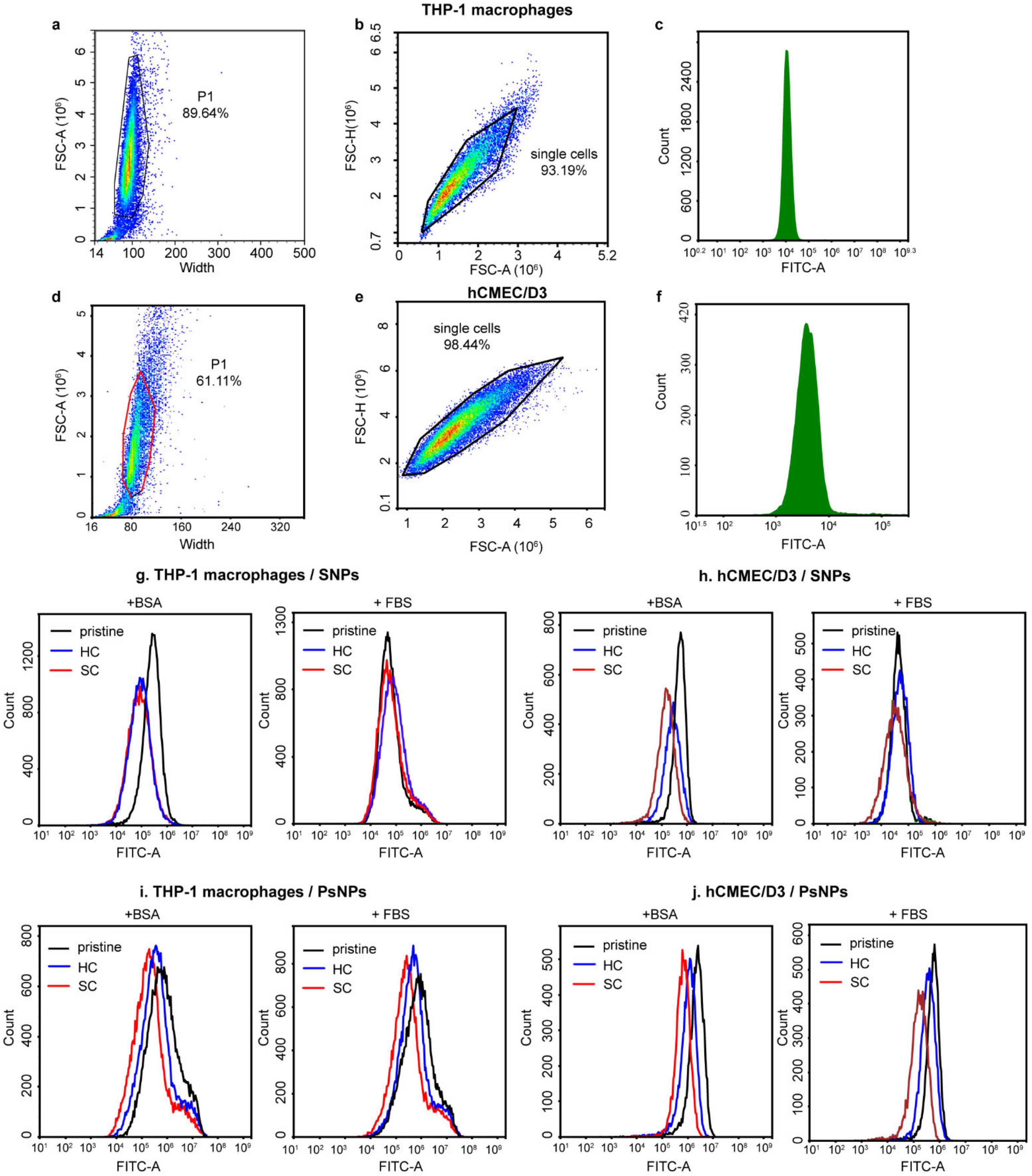


**Supplementary Fig. S16. Flow cytometry-gating strategy for cell association of nanoparticles. a,** THP-1 macrophage cell debris were excluded in a forward scatter/ width dot plot (FSC-A vs width) and the gate was applied to the samples. **b,** A FSC-H vs FSC-A dot plot was used to select THP-1 macrophage single cells. **c**, The median fluorescence intensity of control samples (untreated with nanoparticles) was determined via a histogram. **d-f,** The same strategy as what used for THP-1 macrophage cells was also applied to the hCMEC/D3 cells. **g,h,** The median fluorescence intensity of SNPs associated with THP-1 macrophages (**g**) and hCMEC/D3 cells (**h**) in RPMI media supplemented with BSA or FBS. **i, j**, The median fluorescence intensity of PsNPs associated with THP-1 macrophages (**i**) and hCMEC/D3 cells (**j**) in RPMI media supplemented with BSA or FBS.

**Supplementary Fig. S17.**

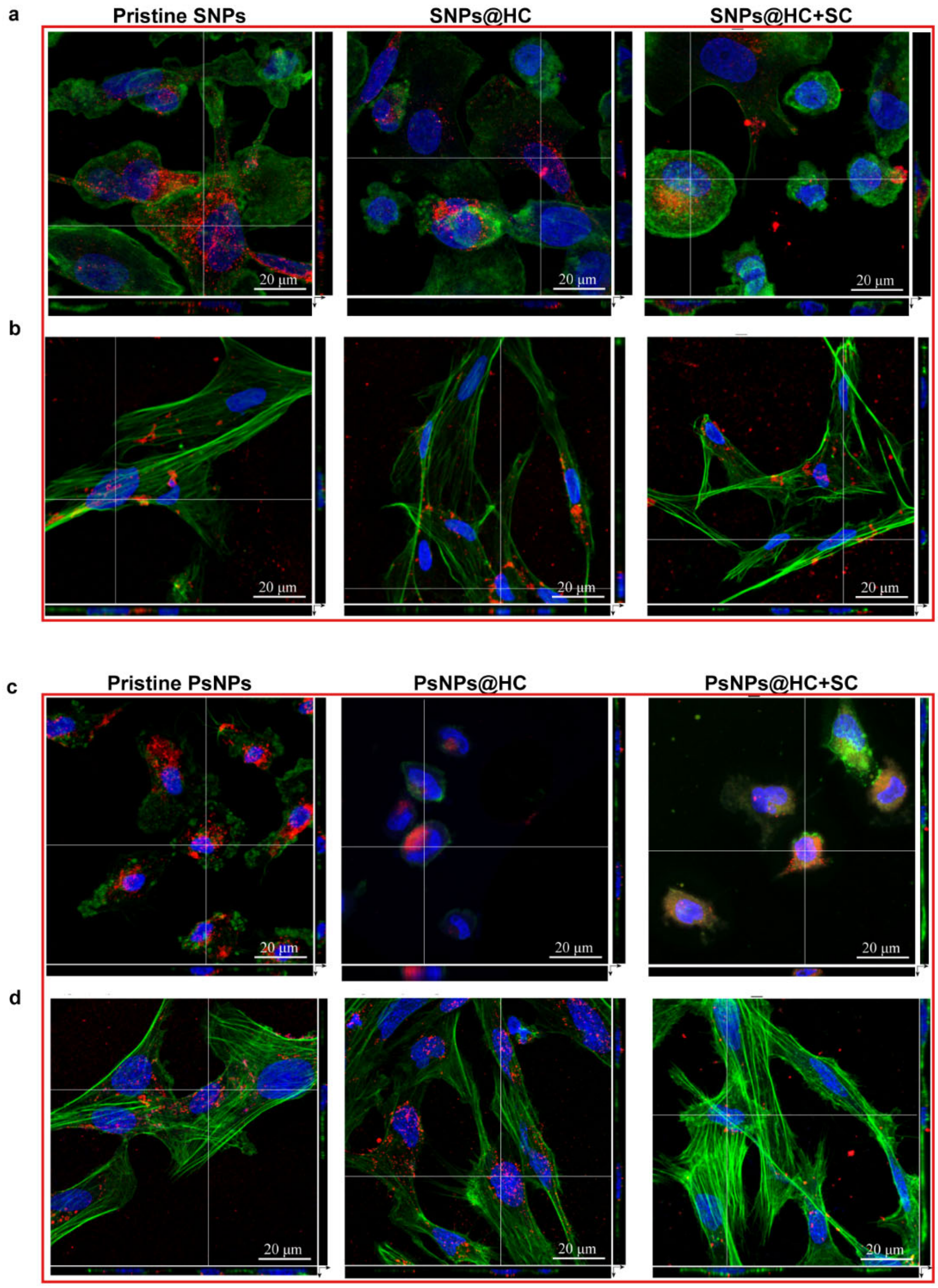


**Supplementary Fig. S17. CLSM confirm the uptake of nanoparticles. a-b,** Orthogonal views of 3D stacks of CLSM images of SNPs–corona complexes confirm the uptake of nanoparticles in THP-1 macrophages (**a**) and hCMEC/D3 cells (**b**) in RPMI containing BSA. **c,d**, Orthogonal views of 3D stacks of CLSM images of PsNPs–corona complexes confirm the uptake of nanoparticles in THP-1 macrophages (**c**) and hCMEC/D3 cells (**d**) in RPMI containing BSA. The images demonstrate that by vigorous washing of nanoparticles, all non-internalized particles had been removed from the cell surface prior to flow cytometry analysis and they are internalized by both cell types. In CLSM images, no differences in the localization of NPs inside the cells were observed. The cells grown on collagen-coated coverslips were fixed and then the cell nuclei (blue) and the actin filaments (green colour) were stained with Hoechst and phalloidin, respectively. The FITC labeled nanoparticles are shown in red colour. Bar, 20 µm.

**Supplementary Fig. S18.**

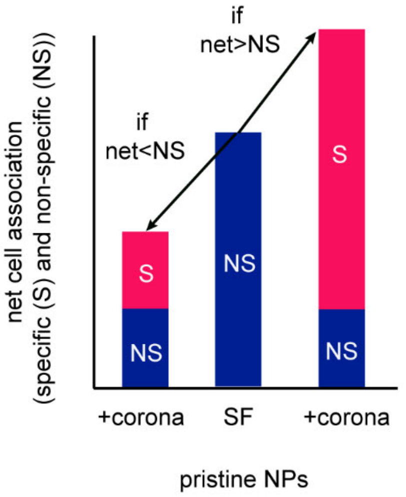


**Supplementary Fig. S18.** The final cell association of the nanoparticle-corona complexes will be determined by the net effect of non-specific (NS) interactions of the particle surface with the cells and specific interactions (S) with the HC, which suggests that the properties of the bare particle surface can still directly influence cell association after a protein corona is formed. Depending on the number of specific interactions made by HC, the final net cell association can be more or less than the non-specific interaction of pristine nanoparticles in the serum-free (SF) medium.

**Supplementary Fig. S19**

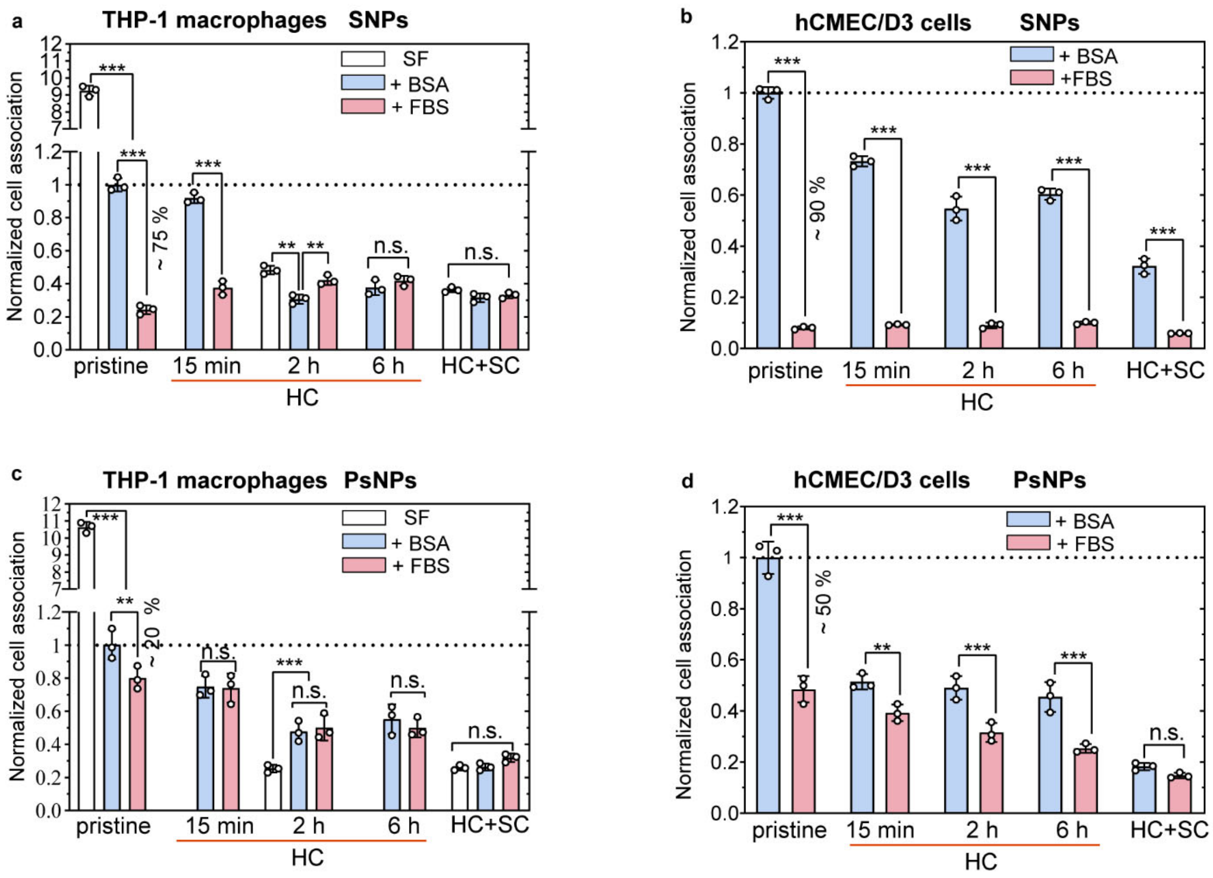


**Supplementary Fig. S19. Comparison of cells association of SNPs-corona complexes and PsNPs-corona complexes in RPMI supplemented with BSA and FBS, shown in Fig.4. a ,b,** Comparison of the cell association of SNPs-corona complexes (**a**) and PsNPs-corona complexes (**b**) in THP-1 macrophages. **c,d,** Comparison of the cell association of SNPs-corona complexes (**c**) and PsNPs-corona complexes (**d**) in hCMEC/D3 cells. The cells were exposed to the pristine NPs, NPs coated with HC formed over different FBS exposure times (15 min, 2 h, and 6 h), and NPs coated with HC_SC for four hours in serum-free RPMI or supplemented with 0.5 % BSA or 10 % FBS. The flow cytometry data were normalised to the pristine nanoparticles values in the RPMI supplemented 0.5 % BSA. Bars shows mean ± sd. of three independent experiments. * p<0.05; ** p< 0.01; *** p<0.001; n.s., not significant.

**Supplementary Fig. S20.**

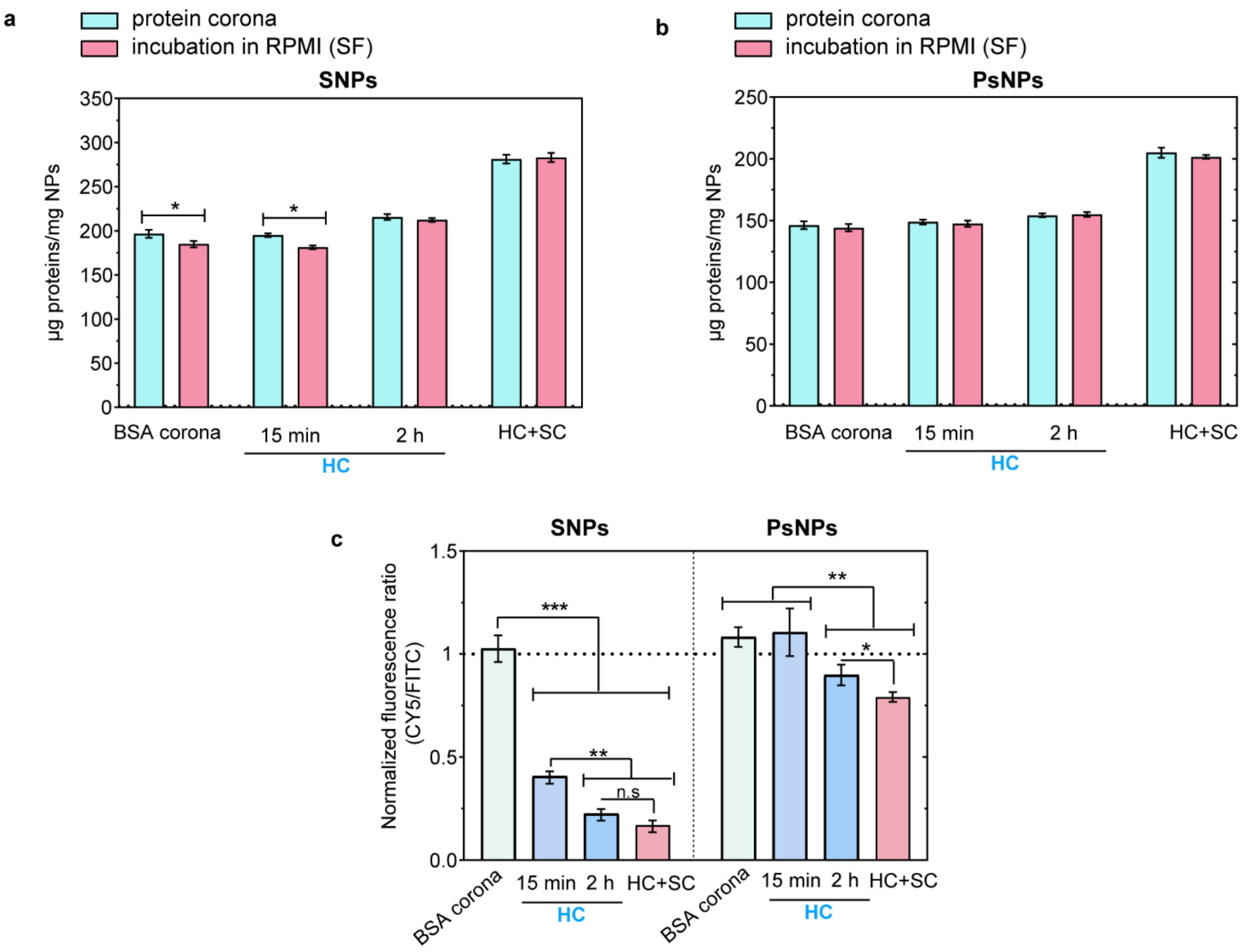


**Supplementary Fig. S20. Stability of proteins on nanoparticles. a,b,** stability of corona proteins on SNPs (**a**) and PsNPs (**b**) in serum-free RPMI. The nanoparticles with BSA corona, FBS corona (HC), and HC-SC were incubated in RPMI (SF) for 2 h. BSA corona and HC_15 min on SNPs were less stable than HC_2h and HC-SC, while all protein coronae on PsNPs were stable. **c,** Exchange of corona proteins on SNPs and PsNPs with CY5 labeled BSA (BSA-CY5). 5 mg/ml BSA-CY5 was added to the nanoparticles with different corona (BSA, HC_15 min, HC_2 h, and HC-SC) and incubated for 2 h. The CY5 fluorescence is representative of BSA proteins exchanged with or added to preformed proteins and FITC fluorescence is for nanoparticles. The fluorescence data were normalized to the BSA corona values on SNPs. Bars show mean ± sd. of three independent experiments. * $p<0.05$; ** $p< 0.01$; *** $p<0.001$;

**Supplementary results**

**Optimization of the click chemistry reaction for capturing weakly interacting proteins**

Sulpho-SASD and DBCO-Sulpho-NHS were used for modification of proteins to perform the click chemistry reaction described in Fig..1a. The modification occurs through the reaction between Sulpho-NHS moieties on the crosslinkers with primary amines on proteins. None of the azide or DBCO reactive groups on these heterobifunctional crosslinkers react with any of the functional groups on proteins, which avoid crosslinking of HC or SC proteins with other HC or SC ones. Moreover, Sulpho-SASD contains a dithiol, which provides a possibility to cleave the covalent bond between proteins by using reducing agents for analysis.

In step 1, HC was first formed through incubation of SNPs with Fetal Bovine Serum proteins (FBS) for 2 h (Fig.1 a, step 1), followed by centrifugation steps to remove weekly bound or unbound proteins from nanoparticles (Fig1a, step 2). Then, the HC proteins on SNPs were modified with different concentrations of Sulpho-SASD (Fig1a, step3). To confirm the labelling, the azide-modified particles were incubated with DBCO-Sulpho-CY5 which reacts with the azide groups through a SPAAC click reaction (Supplementary Fig.1a). The labelling efficiency was measured using two methods. In the first method, the amount of proteins was calculated by a combination of SDS-PAGE and BCA assay. In the second method, the LC-MS/MS data was used to calculate the protein content on SNPs. The results show that the labelling efficiency increases with Sulpho-SASD concentration and reaches a plateau at 0.55 mM Sulpho-SASD. At this concentration, there is at least one aizde on each hard corona protein. Converting $N_3$ group by UV to nitren decreased the click reaction efficiency, which is considered as a control to prove that both $N_3$ and DBCO are necessary for the click reaction.

The fluorescence image and the densitometry analysis of the SDS-PAGE of CY5 labeled proteins eluted form SNPs also confirmed that all HC proteins on SNPs were modified with azide (Supplementary Fig.1b and c). No CY5 fluorescence was detected for the un-labelled corona proteins in the fluorescence image, which indicates that DBCO-CY5 only reacts with N3 modified proteins.

Then, free FBS proteins were modified with DBCO (step 4). The concentration of DBCO (0-0.8 mM) was optimized by measuring the degree of labelling (Supplementary Table.S1). The UV-absorbance labelled proteins are shown in Supplementary Fig. 2a. The DOL increases from 4.2±0.3 to 5.1±0.2 by increasing DBCO-Sulpho-NHS concentration from 0.2 to 0.4, which only increases to 5.6±0.1 when the concentration is 0.8 mM. DOL did not significantly change when proteins were labelled with both CY5 and DBCO (Supplementary Table S1).

Then, DBCO modified proteins were added to SNPs@HC-N3 to capture weakly interacting proteins (Supplementary Fig.2b,c). The SDS-PAGE analysis showed an addition of some proteins by the click reactions. The appearance of some new bands or intensifying of some other bands (especially the one close to 40 kDa and the ones between 50-200 KDa) captured by click reaction is clear in the densitometry analysis (Supplementary Fig.2c). The BCA assay showed that increasing DBCO concentration from 0.4 to 0.8 mM did not lead to a significant change in the amount of corona proteins.

**Supplementary references**


1. Monopoli, M. P. *et al.* Physical- chemical aspects of protein corona: relevance to in vitro and in vivo biological impacts of nanoparticles. *J. Am. Chem. Soc* **133,** 2525–2534 (2011).

2. Tenzer, S. *et al.* Rapid formation of plasma protein corona critically affects nanoparticle pathophysiology. *Nat Nanotechnol* **8,** 772–81 (2013).

3. Fleischer, C. C. & Payne, C. K. Secondary structure of corona proteins determines the cell surface receptors used by nanoparticles. *The Journal of Physical Chemistry B* **118,** 14017–14026 (2014).

4. Yan, Y. *et al.* Differential roles of the protein corona in the cellular uptake of nanoporous polymer particles by monocyte and macrophage cell lines. *ACS nano* **7,** 10960–10970 (2013).
